# Ultra-sensitive FLORA-seq links cell-type-specific tRNAome dynamics to differentiation trajectories guiding therapeutic suppressor tRNA candidate selection

**DOI:** 10.64898/2026.04.14.718460

**Authors:** Hao Li, You-Jia Zhou, Zhen Zhang, Jian-Yang Ge, Zi-Ce Zhou, Wen-Yu Zhu, Xing-Yi Wu, Jiang Wu, Pei-Yu Tian, Yawei Gao, Jianlong Sun, Ru-Juan Liu

## Abstract

Mammalian genomes encode hundreds of tRNA genes, but the role of individual tRNAs in development and cell identity remains unclear. Here, we introduce FLORA-seq, a method for simultaneous, low-input profiling of tRNAs, tRNA-derived RNAs (tdRs) and their modifications from as few as 5-20 cells. Applying FLORA-seq to mouse hematopoiesis revealed 50 high-variance isodecoders whose expression profiles recapitulate known hematopoietic cell types and correlate with differentiation trajectories. At the isoacceptor level, tRNA pools are largely stable within hematopoietic stem and progenitor cells (HSPCs), showing only subtle, specific variations, but become both distinct from HSPCs and internally stable within each terminally differentiated lineage. We further detected dynamic changes in tdR ratios and modification patterns, underscoring precise tRNA regulation during differentiation. Crucially, analysis of 54 endogenous tRNA isodecoders and their engineered suppressor counterparts showed that efficient premature termination codon readthrough preferentially arises from highly expressed cognate isodecoders. This correlation provides a rational framework for prioritizing suppressor tRNA candidates and implies non-redundant functions for individual isodecoders.

## Main

tRNAs are crucial adaptors in protein synthesis and also regulate diverse cellular processes^1–3^. Defects in tRNA function not only contribute to human diseases (e.g., neurological/metabolic disorders and cancer)^1,2,4–6^ but also impair core cellular processes like proliferation and differentiation^3,7^. In mammals, over 600 tRNA genes encode merely 49 isoacceptor families, yet each isoacceptor encompasses multiple isodecoders^8–12^, resulting in tRNAs constituting ∼15% of total RNAs^13^. Notably, the abundance of tRNA isodecoders varies markedly across species, with over 50% of mammalian tRNA genes encoding these variants^10^. This organization has led to the perception of redundancy among tRNA genes. Nevertheless, recent research showed that individual tRNA isodecoders could also have cell-type-specific expression and unique functions^14,15^. tRNA^Arg^-TCT-4-1 was first discovered to exhibit neuron-specific expression patterns with markedly reduced expression levels in glial cells^14,15^. Furthermore, the loss of this particular isodecoder results in functional deficits that cannot be compensated by other isodecoders within the same family^14,15^. These findings demonstrate that different isodecoders may be selectively expressed depending on cell type, developmental stage, or environmental conditions, potentially playing an active role in regulating translation, rather than merely being products of gene duplication and neutral mutation. Studies at the isoacceptor level have established a link between tRNA pool composition and cellular state transitions^3^. Recent work employing ChIP-seq and tRNA-seq has revealed isodecoder-level tRNA expression patterns in defined settings, including distinct neural cell types^11^ and *in vitro* induced differentiation^12^. However, how tRNA repertoires are dynamically remodeled across continuous *in vivo* differentiation processes, particularly within rare progenitor populations, remains poorly understood. Meanwhile, the application of engineered tRNA for therapy is emerging as a promising field^16^, but how endogenous tRNA expression affects therapeutic efficiency and strategies to optimize engineered tRNA design remain unclear.

Furthermore, tRNAs can be processed into tRNA-derived RNAs (tdRs, also known as tRNA-derived fragments or small RNAs, tRFs/tsRNAs) in response to diverse cellular and environmental stimuli, thereby generating a novel class of non-coding RNA regulators that modulate gene expression at multiple levels^2,17–21^. tdRs have been reported to play significant roles in transgenerational inheritance^20,22^, tumorigenesis^21^, aging^23^, development^24^, and differentiation^25^. The abundance of tdRs is dynamically regulated by tissue-specific requirements, cellular states, and environmental conditions^17,18^. Additionally, tRNAs are characterized by over 100 types of chemical modifications, with an average of approximately 13 modifications per mature tRNA^26^. These modifications exhibit tissue– and cell-specificity and can be reprogrammed under stress conditions^1,26–30^. Chemical modifications introduce an additional layer of regulatory complexity to the functions of both tRNAs and tdRs. Systematic exploration is required for tdR expression patterns as well as the dynamic remodeling of tRNA/tdR modifications during developmental cascades. From this integrated perspective, the tRNAome constitutes a dynamic regulatory network encompassing tRNAs, tdRs, and their modifications, which collectively orchestrate cellular functions. Deciphering the dynamics of this integrated tRNAome network, especially at cell-type-specific resolution, requires correspondingly advanced profiling tools.

The multifaceted complexity of tRNA (including diverse genes, intricate structures, and extensive modifications) poses significant sequencing challenges. Consequently, the cell-type-specific distribution patterns of tRNA are not fully understood. NGS-based methods to address these limitations involve the use of RNA demodification enzymes (e.g., AlkB) and/or specialized reverse transcriptases (e.g., TGIRT, MarathonRT) to reduce reverse transcription (RT) termination caused by Watson-Crick face modifications^31–43^. More recent methods have streamlined library preparation and reduced input requirements. mim-tRNAseq^34^ and Induro-tRNAseq^40^ accept total RNA as input and enable detection of a subset of tRNA modifications. MSR-seq further allows co-detection of tdRs with input amounts as low as 10 ng of total RNA^39^. OTTR eliminates conventional ligation steps, thereby simplifying the experimental workflow^35^. To further enable direct profiling of ultra-low-input samples at picogram levels, such as rare hematopoietic stem cells and early embryonic cells, where simultaneous detection of tRNAs and tdRs remains challenging, we established <u>F</u>ast and <u>Lo</u>w-input <u>R</u>NA <u>A</u>tlas sequencing (FLORA-seq). FLORA-seq enables simultaneous detection of full-length tRNAs, tRNA fragments, and modification-sensitive reverse transcription signatures directly from as few as 5-20 cells, without requiring RNA size selection or enzymatic demethylation. FLORA-seq incorporated several key optimizations, such as enhanced poly(A) tailing, utilizing template switching to eliminate adapter ligation, and efficient 3’ end preprocessing for tRNA sequencing. These optimizations improved library construction efficiency, reduced premature RT termination, and allowed for modification detection. FLORA-seq thus provides accurate and reproducible tRNA and tdR profiling from low-input samples, enabling the characterization of rare cell populations and dynamic cellular processes.

Hematopoiesis serves as a paradigm for studying hierarchical cellular differentiation due to its well-defined ontogeny and extensive functional diversity across lineages. Leveraging its ultra-low-input capability, FLORA-seq analyzes rare progenitor populations inaccessible to bulk methods and performs systematic, simultaneous profiling of tRNAs and tdRs. This approach thereby delineates tRNAome regulation from stem/progenitor states through lineage commitment. Our results reveal dynamic changes in tRNAome expression during hematopoietic differentiation. Surprisingly, cell-type-specific isodecoder expression patterns enable robust clustering of 26 hematopoietic populations, highlighting tRNA expression landscapes as an informative layer of cellular identity. Based on the cell-type-specific isodecoder expression, we designed 54 suppressor tRNAs (sup-tRNAs). Their premature termination codon (PTC) readthrough efficiency directly correlates with the expression levels of their cognate endogenous isodecoders. This finding suggests non-redundant functions of individual isodecoders and illustrates the utility of FLORA-seq in prioritizing candidate sup-tRNAs for further therapeutic development.

## Results

### Streamlined FLORA-seq enables user-friendly, ultra-low-input tRNA profiling

tRNAs adopt a cloverleaf-like secondary structure (Fig. 1a) that further folds into a “reverse L”-shaped tertiary structure, making tRNA sequencing particularly challenging. Additionally, chemical modifications on the Watson-Crick face (e.g., m^1^A, m^1^G, m^2^_2_G, m^3^C, acp^3^U) and bulk size modifications (e.g., t^6^A, mcm^5^s^2^U, Fig. 1a, b) can obstruct RT, leading to shortened cDNA synthesis^33^. Existing methods that aim to address these challenges require cumbersome steps, making it difficult to detect tRNAs in ultra-low RNA input. To overcome these limitations and enable tRNA profiling from ultra-low RNA inputs, we optimized the library preparation workflow. Key refinements included: selecting a robust reverse transcriptase, adopting an efficient ligation strategy, implementing template switching to replace a second ligation/circularization step, and designing custom adapters. These were integrated into a streamlined, all-in-one process for tRNA library construction.

**Fig. 1.**
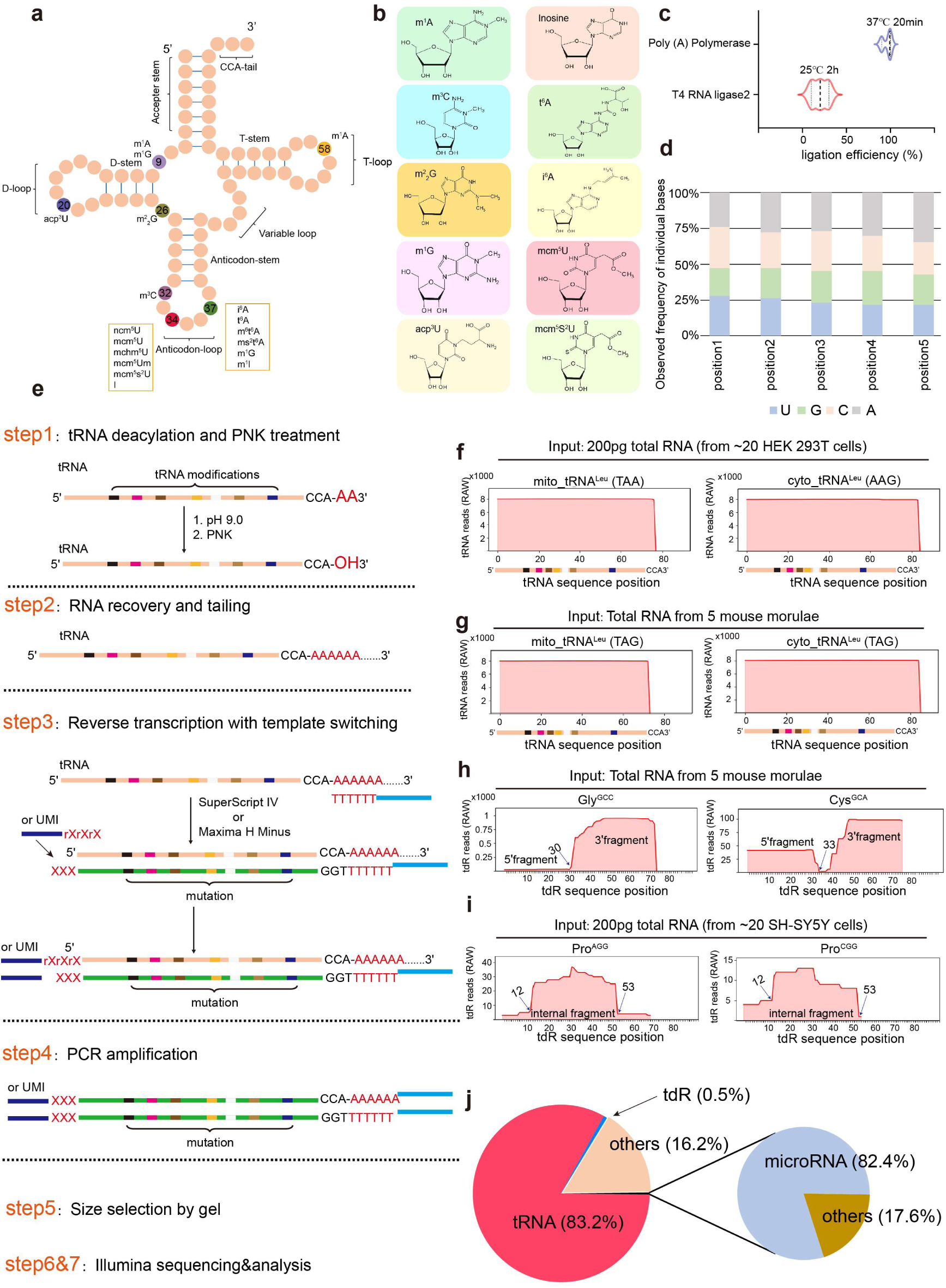
The FLORA-seq workflow. **a,** Cloverleaf structure of tRNA with some modifications. m^1^A, 1-methyladenosine; m^1^G, 1-methylguanosine; acp^3^U, 3-(3-amino-3-carboxypropyl) uridine; m^2^_2_G, *N*^2^,*N*^2^-dimethylguanosine; m^3^C, 3-methylcytidine; ncm^5^U, 5-carbamoylmethyluridine; mcm^5^U, 5-methoxycarbonylmethyluridine; mcm^5^s^2^U, 5-methoxycarbonylmethyl-2-thiouridine; I, inosine; i^6^A, *N*^6^-isopentenyladenosine; t^6^A, *N*^6^-threonylcarbamoyladenosine; m^6^t^6^A, *N*^6^-methyl-*N*^6^-threonylcarbamoyladenosine; ms^2^t^6^A, 2-methylthio-*N*^6^-threonylcarbamoyladenosine; m^1^I, 1-methylinosine. **b,** Chemical structures of some tRNA modifications. **c,** The ligation efficiency of Poly(A) polymerase and T4 RNA ligase 2. Using an equal amount of total tRNA without any adapter ligation or poly(A) addition as a control, the ligation efficiency is calculated as the ratio of RNA molecules with successfully added adapters or poly(A) tails to those in the control. **d,** A pool of RNA oligonucleotides with identical sequences but varying 5’-terminal nucleotides (A, C, G, or U) was subjected to the template-switching reverse transcription library preparation under optimized conditions. The bar graph shows the proportion of each oligonucleotide recovered in the final sequencing libraries. **e,** Schematic overview illustrating the steps required for tRNA library preparation using FLORA-seq. **f,** Raw read coverage of mito_tRNA^Leu^ (TAA) and cyto_tRNA^Leu^ (AAG) from 20 HEK 293T. **g,** Raw read coverage of mito_tRNA^Leu^ (TAG) and cyto_tRNA^Leu^ (TAG) from 5 mouse morulae. **h,** Raw read coverage of 3’ and 5’tdR from 5 mouse morulae. **i,** Raw read coverage of internal tdRs from 5 mouse morulae. **j,** Pie chart illustrating the distribution of distinct RNA species identified from non-ribosomal genome-aligned reads in mouse morula sequencing data. Reads with mapped lengths below 15 bp were excluded from the analysis.

To optimize RT enzyme performance and reaction conditions, we evaluated four RT enzymes known for their ability to tolerate modifications and induce mutations: HIV RT mutant RT1306^42^, MarathonRT^36,43^, SuperScript IV^37–39^ and Maxima H Minus^44,45^. Firstly, RT was performed on synthetic and endogenous tRNA using four RT enzymes (Extended Data Fig. 1a-c). Next, we designed primers at the 3’ end of the cDNA and used qPCR to detect full-length cDNA (Extended Data Fig. 1a-c). When using synthetic tRNA as the template, all four enzymes generated comparable yields of full-length cDNA (Extended Data Fig. 1b, left). Using endogenous tRNAs as template, SuperScript IV and Maxima H Minus exhibited better capability in generating full-length cDNA (Extended Data Fig. 1b, right). Their comparable yields from synthetic and endogenous templates (Extended Data Fig. 1c) demonstrated that SuperScript IV and Maxima H Minus overcome modifications without compromising RT efficiency. Based on these results and Maxima H Minus’s reported ability in handling complex RNA modifications^44^, we selected it for our experiments.

Additionally, we optimized the temperature and time of RT for ultra-low input. It has been reported that prolonged incubation under optimal temperature is beneficial for the synthesis of full-length cDNA^34,40^. Therefore, we tested the cDNA synthesis capability of Maxima H Minus at low (50°C) and higher temperatures (60°C) for 2, 3, and 12 h, respectively. The results showed that when using synthetic tRNA as a template, neither temperature nor time affected the synthesis of cDNA (Extended Data Fig. 1d), indicating that the synthetic tRNA is suitable as an internal control. For endogenous tRNA, the full-length cDNA yield at 60°C was 30% lower than at 50°C, and extending the reaction to 12 hours did not mitigate this difference (Extended Data Fig. 1e). Conversely, at 50°C, yields after 2 or 3 hours were approximately 20% higher than after 12 hours (Extended Data Fig. 1e). The increase in cDNA yield at 50°C, rather than at 60°C, may be due to reduced dissociation of the RT enzyme from the tRNA templates at the lower temperature^40^. Thus, to generate sufficient cDNA, a 2-3 h RT period at 50°C is considered optimal.

To distinguish mature tRNAs from pre-tRNAs, we tested two approaches: ligating adapters or elongating a poly(A) tail at the 3’ end of tRNA. Since over 80% of tRNAs are aminoacylated at 3’ ends^46^, we first deacylated the samples to expose the 3’ hydroxyl group, enabling the addition of DNA adapters or poly(A) tails. Subsequently, we compared the efficiency of adapter ligation with that of poly(A) tailing. The results indicated that only approximately 30% of tRNA could be ligated with adapters within 2 h (Fig. 1c and Extended Data Fig. 1f), whereas poly(A) tails were successfully added to nearly all tRNAs within 20 min (Fig. 1c and Extended Data Fig. 1f). Therefore, we adopted poly(A) tail elongation at the 3’ end of tRNAs.

For streamlined experimentation, we implemented template switching, allowing the concurrent introduction of a known sequence at the 3’ end of cDNA during RT. This strategy eliminates the need for separate 5’ adapter ligation or cDNA circularization steps, thereby simplifying the workflow and making it particularly robust for ultra-low-input samples. To evaluate potential sequence bias in template switch oligo (TSO) addition, we performed a ligation efficiency assay using four RNA oligonucleotides with different 5’-end nucleotides (X = A, C, G, U; N5X-3’) as input samples. Following TSO addition, the resulting cDNA was amplified and subjected to next-generation sequencing. The results showed that under optimized reaction conditions, all four RNA oligonucleotides generated stoichiometric amounts of ligation products (Fig. 1d), indicating no obvious bias in TSO addition toward any specific 5’-end nucleotide. Finally, to achieve an “all-in-one” approach, we utilized TSO and Oligo(dT) pre-designed with specific sequences, enabling direct initiation of library amplification. Thus, the template-switching activity of Maxima H Minus, combined with sequence-specific primers, enabled direct post-RT cDNA amplification.

Building on these insights, we established FLORA-seq, a streamlined process for ultra-low input tRNA sequencing (Fig. 1e). Using this approach, we successfully profiled tRNAs and tdRs from as few as 5-20 HEK 293T cells or 5 mouse morulae (Fig. 1f-1j, Extended Data Fig. 1g-k, and Supplementary Tables 1-3), demonstrating its sensitivity.

### FLORA-seq enables integrated analysis of tRNAs and tdRs

To evaluate the sensitivity of FLORA-seq, we tested a range of total RNA inputs from 200 pg to 20 ng, corresponding to approximately 20 to 2,000 HEK 293T cells, prepared by serial dilution from a single 100 ng stock. To further demonstrate applicability to intact low-input samples, we successfully generated FLORA-seq libraries from RNA purified from 5 HEK 293T cells and from 5 mouse morulae. Although libraries were successfully generated from 5 cells, we excluded this extreme low-input condition from downstream analysis owing to the presence of adapter self-ligation artifacts. Despite using very small amounts of total RNA as input, FLORA-seq successfully mapped over 50% of reads to tRNAs among all reads mapped to genome (Extended Data Fig. 2a), demonstrating high efficiency in library preparation. We identified 203 unique tRNA isodecoders in HEK 293T cells (Supplementary Table 1) and 177 in mouse morulae (Supplementary Table 2). Our method achieved comparable tRNA isodecoder identification to high-input methods (e.g., mim-tRNAseq detecting 216 isodecoders) with only 5-20 HEK 293T cells, demonstrating that FLORA-seq can resolve complex tRNA transcriptomes from ultra-low inputs, representing a major step towards genuine single-cell tRNA sequencing. To confirm the capture of mature tRNAs, we analyzed the 3’ ends of uniquely mapped reads. We observed that over 97% of reads ended with “CC”, consistent with the removal of the 3’ terminal adenine (A) during adapter trimming (Extended Data Fig. 2b, c). These results indicate that the majority of reads captured by FLORA-seq correspond to mature tRNAs, even under ultra-low RNA input conditions. Moreover, over 90% of uniquely mapped reads were full-length tRNA transcripts (Fig. 2a, Extended Data Fig. 3), showing stable end-to-end coverage for tRNAs (Extended Data Fig. 2d), collectively confirming that optimization prevented truncation and ensured efficient RT with intact tRNA. Further analysis at the isodecoder level also revealed strong agreement between the high (∼20 ng) and low (∼200 pg) input amounts, demonstrating the robustness of FLORA-seq (Extended Data Fig. 2e).

**Fig. 2.**
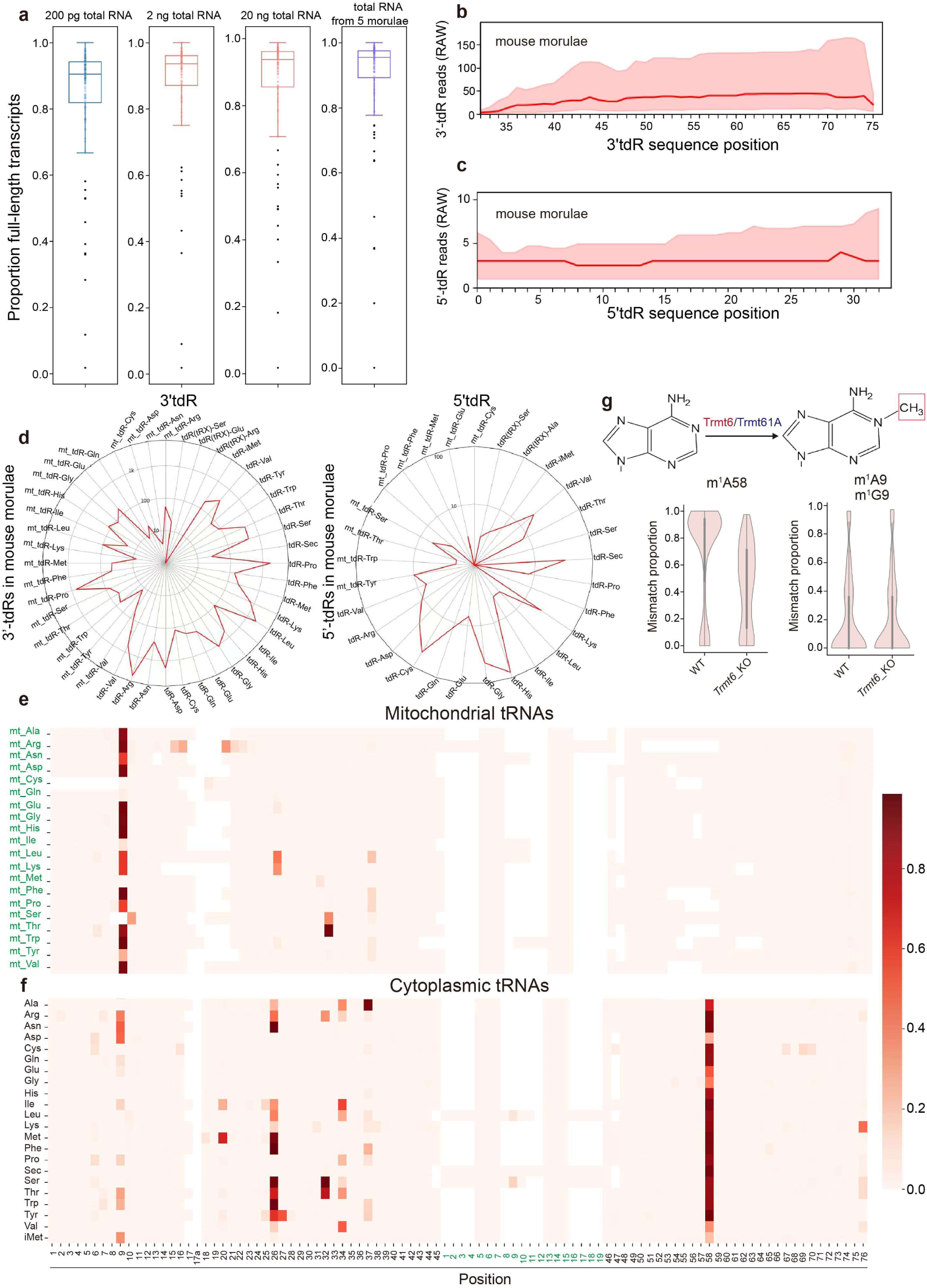
Detect tRNA, tdR, and their modifications simultaneously in ultra-low input samples using FLORA-seq. **a**, Proportion of full-length tRNAs (defined as mapped length > 95% of the total isodecoder transcript length) across samples. Each dot represents one unique tRNA isodecoder. Centerline indicates the median; box limits represent the upper and lower quartiles, outliers shown as black dots. **b, c,** Overall length mapping showing the distribution of relative tdR reads from mature tRNAs presents, for mouse morulae, the median (red line) and interquartile range (pink area) of read numbers for 3’tdRs (**b**) and 5’tdRs (**c**) across different isodecoders. **d,** Distribution of 3’tdRs (left) and 5’tdRs (right) originating from different isodecoders in mouse morulae. Bars represent raw counts displayed on a logarithmic scale. **e, f,** Heatmap showing the mean mismatch level at each base of mitochondrial (**e**) and cytoplasmic (**f**) tRNAs. For each isoacceptor (y-axis), the value represents the mean value of mutation ratio per base across all its tRNA transcript length. **g,** Structure of the m^1^A modification at position 58 of tRNA catalyzed by Trmt6-Trmt61A (upper). Violin plot of mismatch levels at the m^1^A site in wild-type mouse morulae and *Trmt6* knockout morulae (lower). The plot illustrates the distribution of mismatch levels on m^1^A site across all isodecoders.

To comprehensively evaluate the accuracy of FLORA-seq, we benchmarked it against established tRNA-seq methods. Using matched samples, FLORA-seq showed high concordance with Induro-tRNAseq^40^ (Pearson r = 0.68-0.80 across two cell lines), and with mim-tRNAseq^34^ (Pearson r = 0.67) (Extended Data Fig. 2f, g). To further validate the accuracy of FLORA-seq at the isodecoder level, we employed a non-sequencing experimental approach for cross-verification. This method is based on the 5’→3’ exonuclease activity of Taq DNA polymerase, which efficiently hydrolyzes probes only when they perfectly match the template^47^. Based on this principle, we designed TaqMan probes and PCR primers for six tRNA^Arg^-TCG isodecoders (tRNA^Arg^-TCG-1 through tRNA^Arg^-TCG-6). Using Maxima H Minus enzyme for RT on RNA either treated or untreated with AlkB, we confirmed that the expression levels of tRNA^Arg^-TCG isodecoders were highly consistent with the sequencing results (Extended Data Fig. 2h, i). These results demonstrate the reliability and high sensitivity of FLORA-seq.

tdRs are derived from tRNAs and are essential in diverse biological processes^1,20^. However, ultra-low RNA input poses a significant challenge for the simultaneous and robust profiling of both tRNAs and tdRs. tdRs can inherit modifications from their parental tRNAs^48^. Additionally, 3’tdRs typically feature aminoacylated terminal ends, whereas 5’tdRs often contain 3’-P or 2’,3’-cP ^48^. These characteristics complicate adapter ligation. To overcome this, we deacylated the RNA samples and treated them with PNK, which enabled the successful detection of both 3’tdRs and 5’tdRs alongside full-length tRNAs (Fig. 2b, c). Notably, FLORA-seq leverages the template switching activity at the completion of RT, whereby a template-switch oligo is incorporated immediately upon reaching the 5’ terminal of tRNAs or tdRs, ensuring that no RT-stopped sequences are captured. In HEK 293T cells, we observed 69 types of 3’tdRs, while in mouse morula, we observed 74 3’tdRs (Supplementary Table 3). However, internal tRNA derived RNAs (i-tdRs) were virtually undetectable in HEK 293T cells and mouse morulae. Interestingly, we did detect two i-tdRs (i-tdR-Pro-AGG and i-tdR-Pro-CGG) in human neuroblastoma SH-SY5Y cells (Fig. 1i), though their expression remained substantially lower than that of 3’tdR or 5’tdR. Moreover, our results showed that 3’tdR-Arg and 5’tdR-His, which correspond to tRNA halves, were the most abundant tdRs in mouse morulae (Fig. 2d). Notably, we discovered unexpected cleavage of selenocysteine-biosynthesis-linked tRNA^Sec^, directly linking its fragmentation to tdR biogenesis (Fig. 2d).

Together, FLORA-seq provides an efficient and reproducible platform for ultra-low-input co-profiling of tRNAs and tdRs, allowing the characterization of tRNA dynamics in rare cell populations under developmental and disease contexts.

### FLORA-seq captures tRNA and tdR modifications via mutation signatures

tRNA modifications are essential for protein synthesis and cellular function. Their defects or dysregulation are often associated with human diseases^26,27,49^. To determine whether FLORA-seq is sensitive enough to detect different tRNA modifications, we quantified the frequency of base mutations at specific tRNA positions induced by modifications that disrupt Watson-Crick base pairing or impose substantial steric constraints, thereby estimating modification levels. Based on this, we examined the modification profiles of all tRNA isodecoders in both mouse morulae and human cells. We observed that approximately 80% of cytoplasmic tRNAs (cyto_tRNAs) in both species carried the m^1^A58 modification, whereas, mitochondrial tRNAs (mito_tRNAs) predominantly featured m^1^A9 (Fig. 2e, f and Extended Data Fig. 4, 5a). Importantly, the acp^3^U modification at position 20, an attachment site for N-glycans in glycoRNA^50,51^, was also detected (Fig. 2e, f and Extended Data Fig. 4, 5a). In human cells, acp^3^U20 was abundant in tRNAs such as tRNA^Ala^-AGC, tRNA^Asn^-GTT, tRNA^Met^-CAT, tRNA^Thr^-TGT, and tRNA^Thr^-AGT (Extended Data Fig. 4). In contrast, in mouse morulae, acp^3^U20 was highly abundant in tRNA^Met^-CAT and tRNA^Ile^-AAT (Extended Data Fig. 5a). Additionally, we detected bulky modifications in the tRNA anticodon loop, which are critical for translation accuracy and wobble decoding^52–54^. We observed the accumulation of mismatches at positions 32, 34, and 37 (Fig. 2e, f and Extended Data Fig. 4, 5a), confirming these positions were full of modifications as previously reported^1,2^. FLORA-seq, similar to other sequencing methods, identified these modifications with high sensitivity but could not distinguish between modification types present at the same position, such as t^6^A37 and ms^2^t^6^A37.

To validate the accuracy of FLORA-seq in detecting modifications, we focused on m^1^A, one of the most abundant tRNA modifications. We conditionally knocked out *Trmt6*, the enzyme responsible for adding m^1^A58^55–58^, in mouse morulae. We observed a reduction in m^1^A58 level while m^1^A9 remained unchanged. (Fig. 2g and Extended Data Fig. 5a, b). Interestingly, our result showed that m^1^A58 is present on the vast majority of cytoplasmic tRNAs, including tRNA^Gly^-CCC, while mouse tRNA^Gly^-GCC lacks this modification (Extended Data Fig. 5a). To confirm this observation, we designed specific probes for mouse tRNA^Gly^-GCC and tRNA^Gly^-CCC, isolated the endogenous tRNAs from mouse myoblast (C2C12) cell line, and performed RNA mass spectrometry analysis. The results showed high-level m^1^A modification in mouse tRNA^Gly^-CCC, whereas no m^1^A was detected in mouse tRNA^Gly^-GCC (Extended Data Fig. 2j). Together, these findings demonstrate that FLORA-seq can reliably detect specific tRNA modifications.

Furthermore, FLORA-seq enabled the detection of modifications in both 5’tdRs and 3’tdRs (Extended Data Fig. 5c). We observed that tdRs largely inherit modifications from their parental tRNAs, with m^1^A being the most conserved across both types of fragments (Extended Data Fig. 5c), supporting the idea that most tdRs originate from mature tRNAs. In conclusion, FLORA-seq can detect reverse transcription signatures indicative of modifications at specific tRNA positions (9, 20, 26, 32, 34, 37, and 58) and on tdRs, though it cannot distinguish different modification types at the same position nor detect modifications that do not induce RT arrest or misincorporation.

### High-variance tRNA isodecoders clustering maps cell-type identity networks and lineage-specific differentiation trajectories

Cellular differentiation is governed by multilayered regulatory networks, yet whether dynamic regulation of tRNA isodecoders contributes to cell-state specification remains unclear, particularly whether cell-type-specific isodecoders perform nonredundant functions under different cellular contexts. To address this question, we leveraged hematopoiesis, a canonical model of stem cell differentiation into functionally specialized lineages^25,59–66^.

We systematically profiled 26 hematopoietic cell types, from hematopoietic stem cells (HSCs), multipotent progenitors (MPPs, MPP1-MPP6), and lineage-committed progenitors (common myeloid progenitor cells, CMPs; Granulocyte-monocyte progenitor cells, GMPs; megakaryocyte-erythroid progenitor cells, MEPs; common lymphoid progenitors, CLPs) to terminally differentiated immune cells (NK, B, T cells) and blood components (erythrocytes, megakaryocytes), using FLORA-seq to map mature tRNA abundance. This continuum captures key transitions from stemness maintenance to effector function acquisition (Fig. 3a and Extended Data Fig. 6).

**Fig. 3.**
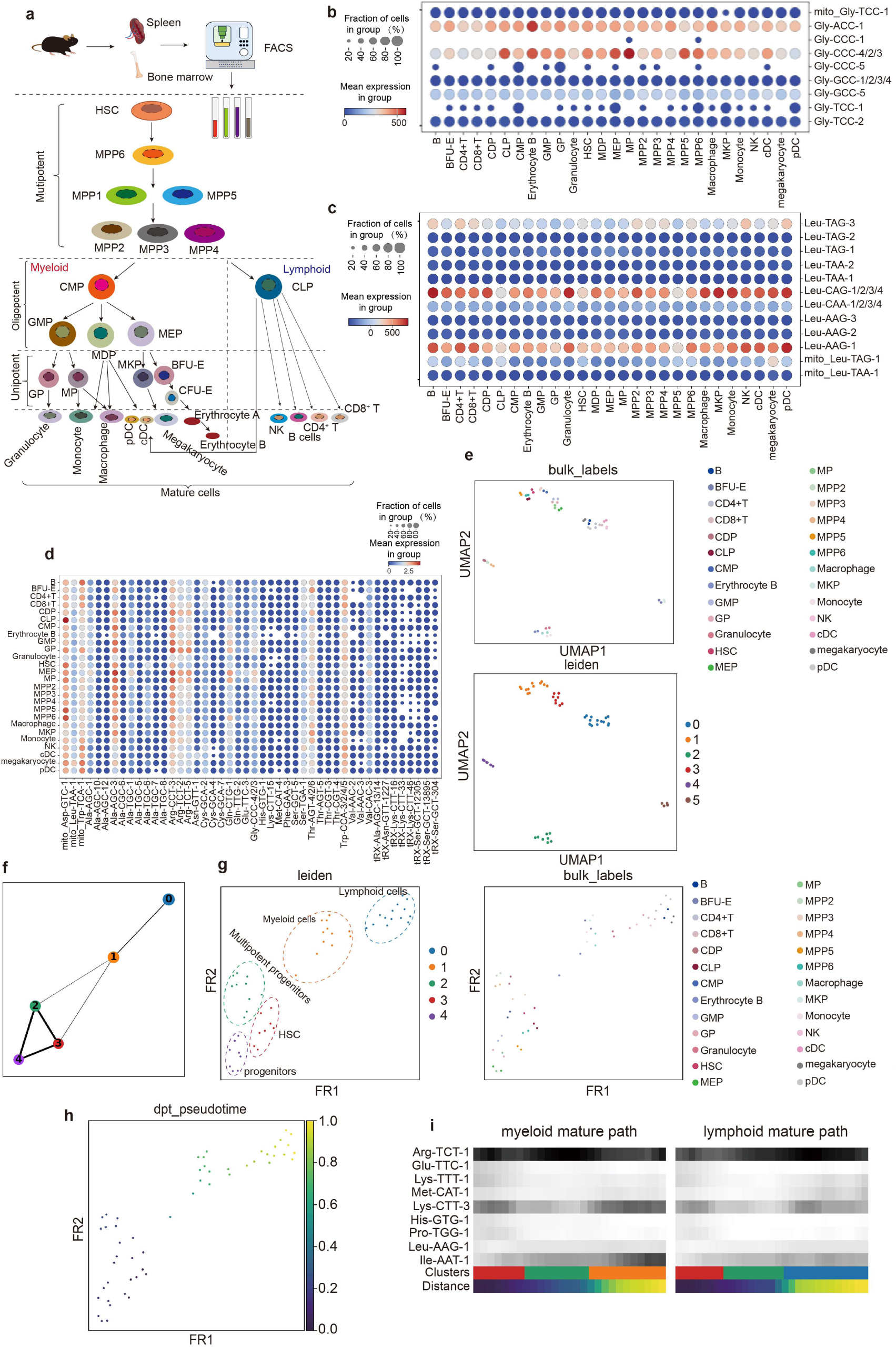
Cell stage-specific characterization through single tRNA isodecoder signatures. **a**, Schematic showing the collection of samples from mouse spleen and bone marrow, followed by cell sorting using FACS markers. The diagram illustrates that cells from 30 hematopoietic populations were targeted, and 26 of them were successfully enriched and applied in downstream experiments. **b, c,** Relative expression levels of tRNA^Gly^ (**b**) and tRNA^Leu^ (**c**) isodecoders across 26 hematopoietic cell types. **d,** Relative expression levels of high-variance tRNA isodecoders across 26 hematopoietic cell types. In (**b-d)**, dot size indicates the reproducibility of isodecoder detection across biological replicates: larger dots correspond to expression in both replicates, while smaller dots indicate expression in only one. Two biological replicates per cell type. **e,** UMAP embedding of cell clusters derived from FLORA-seq across 26 different cell types. Each colored dot represents a cell type, with two biological replicates per type. Leiden clustering result of 5 independent clusters (n_neighbors = 4). **f, g,** Leiden clustering (**f**) and trajectory inference (**g**) of 26 cell types. **f,** Abstracted lineage graph showing inferred connectivity between clusters based on tRNA expression profiles. Nodes represent Leiden clusters results. **g,** Two-dimensional embedding (FR1–FR2) of samples colored by group annotations. Clusters showing major cell types in each group, each dot represents one sample, cell type as indicated in the legend (n_neighbors = 5). **h,** Pseudotime ordering of cells, with the root specified as cluster containing HSCs. The color gradient represents pseudotime progression, illustrating the timeline of hematopoietic development. **i,** Heatmaps showing normalized tRNA isodecoder expression levels during myeloid and lymphoid cell differentiation. Clusters correspond to those in (**e**), with distances based on pseudotime calculations from (**f**).

Strikingly, our high-resolution mapping unveiled pervasive yet previously uncharted isodecoder expression plasticity across differentiation trajectories. We observed that tRNA^Gly^-ACC-1 and tRNA^Gly^-CCC-2/3/4 emerged as dominant isodecoders during hematopoiesis. The expression of these isodecoders differ between cell types, for example, tRNA^Gly^-CCC-4/2/3 showing the highest expression levels in progenitor cells (Fig. 3b). Similarly, even though tRNA^Leu^-CAG-1/2/3/4 and tRNA^Leu^-TAG-3 are the most abundant isodecoders in their isoacceptor group, their expression varies across cell types (Fig. 3c). Specifically, tRNA^Leu^-CAG-1/2/3/4 and tRNA^Leu^-TAG-3 exhibit reciprocal expression patterns, where cell types with high tRNA^Leu^-CAG-1/2/3/4 expression show minimal levels of tRNA^Leu^-TAG-3 (Fig. 3c). Similarly, other anticodons also have one or a few dominant isodecoders, and these isodecoders display cell-type-specific differential expression (Extended Data Fig. 7).

Previous studies show individual isodecoder like tRNA^Arg^-TCT-4-1 have cell-type-specific expression in neurons, and its function cannot be replaced by other isodecoders^14,15^. Here we identified over 200 isodecoders in 26 cell types, and at least 50 isodecoders can be identified as high-variance isodecoders (defined based on normalized expression dispersion; see Methods) (Supplementary Table 4). Although the functional implications of these expression dynamics remain to be fully elucidated, our findings suggest that tRNA isodecoder plasticity constitutes a previously underappreciated regulatory layer in cellular differentiation, extending the role of tRNAs beyond a uniform translational pool. To explore whether tRNA isodecoders expression pattern can provide insights into cell lineage, we built a model linking tRNA isodecoders dynamics pattern to cell types. High-variance tRNA isodecoders were identified using Scanpy, yielding 50 isodecoders which are expressed in at least 10 samples. A color-coded dot plot of expression levels revealed substantial variation in expression among both isodecoders and cell types (Fig. 3d), despite most high-variance tRNA isodecoders being expressed in almost all cell types, the expression of each isodecoder still varies a lot between cell types. Next, we characterized cell types based on their tRNA transcriptome signatures. Filtering isodecoders expressed in more than two cell types, we applied principal component analysis (PCA) and k-nearest neighbor (KNN) analysis to 157 tRNA isodecoder features, then clustering cells with Leiden algorithm. Unsupervised clustering based on tRNA isodecoder expression separated the 26 pre-defined hematopoietic populations into six groups that largely correspond to known stages of differentiation (Fig. 3e). HSCs and early progenitors clustered together, while mature lymphoid cells and dendritic cells formed distinct groups (Fig. 3e).

Partition-based graph abstraction (PAGA) analysis revealed connectivity patterns consistent with known hematopoietic relationships. Cells were grouped into five clusters using Leiden clustering with a bigger group size to facilitate visualization of inter-cluster relationships. We observed HSCs and progenitors showing close relationships to myeloid progenitors, but weaker connections to mature myeloid and lymphoid cells (Fig. 3f, g). Aligning these stages to a common pseudotime axis, with HSCs set as the root, revealed clear progression patterns (Fig. 3h). Certain isodecoders, such as tRNA^Lys^-CTT-3, displayed bursts of expression in specific cell types, while others, like tRNA^Ile^-AAT-1, showed gradual increases during differentiation, particularly in the lymphoid maturation pathway (Fig. 3i). These findings indicate that tRNA isodecoder expression varies across differentiation-associated states and exhibits stage-associated dynamics across intermediate populations.

In summary, our high-resolution mapping reveals 50 high-variance tRNA isodecoders whose expression is both dynamic and exquisitely context-dependent, thereby highlighting an underappreciated layer of tRNA regulation in differentiation. Consistent with cell-type-restricted tRNA isodecoder expression documented in the nervous system using Pol III ChIP-seq^11^, our continuous hematopoietic atlas, by directly quantifying mature tRNA abundance, demonstrates that isodecoder expression can recapitulate the known hematopoietic hierarchy and distinguish cellular identities.

### Stability and divergence of the tRNA anticodon pool across hematopoietic differentiation

Using FLORA-seq, we mapped the dynamic expression of tRNA isodecoders across hematopoietic differentiation, uncovering distinct cell-type-specific programs. In contrast, at the isoacceptor level, *in vitro* differentiation experiments have indicated that the overall tRNA anticodon pool remains relatively stable^12^. However, it was unclear whether this stability is maintained during multi-stage, continuous differentiation *in vivo*. To resolve this, we analyzed the abundance of isoacceptor (or equivalently, the anticodon pool) to examine whether these tRNA pools align with differentiation stages. Clustering of cytoplasmic tRNA isoacceptor expression (Z-score normalized expression across 26 hematopoietic cell types) clearly separated stem/progenitor cells from terminally differentiated cells (Fig. 4a). This pattern suggests that while the anticodon pool remains relatively uniform within hematopoietic stem and progenitor cells (HSPCs) and also within terminally differentiated lineages, it differs markedly between these two broad cellular compartments (Fig. 4a). Consistent with the model of anticodon buffering during differentiation^12^, our analysis of the continuous hematopoietic hierarchy shows that this stability is maintained within both the stem/progenitor compartment and terminally differentiated lineages, notably, the pool compositions differ between these two cellular states. Prior work has also shown that cells switch tRNA expression programs when shifting from proliferation to differentiation, and that mRNA profile changes are coordinated with tRNA pool adjustments to support efficient translation of stage-specific genes^3,7^. These findings together suggest that HSPCs and terminally differentiated cells may have different translation patterns. To further explore this, we analyzed polysome-seq data^66^ by calculating polysome-associated codon usage based on the CDS regions of both polysome-enriched mRNA and total mRNA. The results showed highly similar codon usage between HSCs and monocyte progenitors (MPs) cells (Fig. 4b). FLORA-seq also indicated that the two cell types share similar anticodon pool (Fig. 4a). Together, these findings suggest that HSCs and MPs, as representative HSPCs, may operate within a coherent translational framework. The distinct tRNA isoacceptor pools we observed (Fig. 4a) likely reflect a fundamental reprogramming of the translational substrate between the HSPC compartment and terminally differentiated lineages. Notably, within this overall compartmental stability, individual tRNA isoacceptors exhibited dynamic expression during differentiation. For example, in early progenitors, tRNA^Glu^-TTC progressively decreased from HSCs to MPP6 and MPP5, while tRNA^Tyr^-GTA showed the inverse pattern, rising from HSCs to MPP5 (Fig. 4a). Similarly, terminally differentiated myeloid cells displayed lineage-specific isoacceptor preferences: granulocytes highly expressed tRNA^Lys^-TTT, whereas monocytes and macrophages (sharing the GMP progenitor) showed minimal levels (Fig. 4a). Together, these expression patterns suggest that selective changes in tRNA isoacceptors abundance may accompany translational adjustments during terminal myeloid differentiation, despite overall stability of the anticodon pool.

**Fig. 4.**
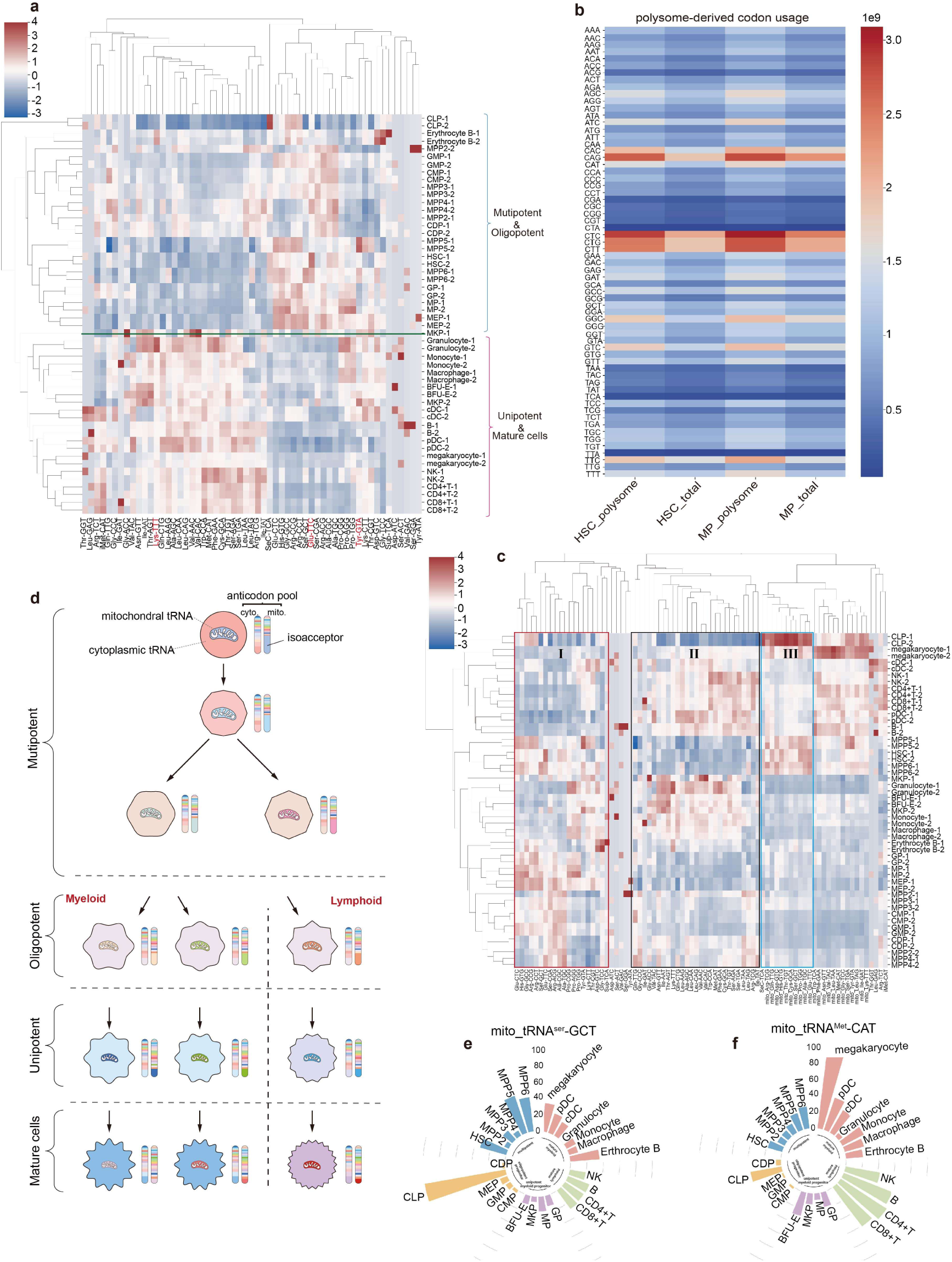
Reprogramming of tRNA isoacceptor landscapes during hematopoietic differentiation. **a**, Clustered heatmap of cytoplasmic tRNA isoacceptors expression across 26 cell types (two biological replicates per type). Expression was normalized to 10,000 counts per sample and further Z-score normalized across samples (values clipped between -4 and 4). Statistical comparisons were performed on normalized raw count data prior to Z-score transformation. A significant difference was observed between HSC and MPP5 for tRNA^Tyr^-GTA (two-sided t-test, *p* = 0.00479, n = 2). tRNA^Lys^-TTT expression differed significantly among granulocytes, monocytes, and macrophages, with higher levels in granulocytes compared to monocytes (*p* = 0.0197, n = 2) and macrophages (*p* = 0.0127, n = 2). **b,** Heatmap of codon usage in HSCs and MPs, based on polysome-enriched RNA and total RNA. Codon usage was calculated by weighting CDS expression levels per gene • codon distribution per gene. **c,** Clustered heatmap of combined cytoplasmic and mitochondrial tRNA isoacceptors expression across 26 cell types (two biological replicates per type). Three tRNA subsets are observed. **d,** Schematic illustration of tRNA pool stability and individual tRNA isoacceptor reprogramming during differentiation. **e, f,** Expression levels of mito_tRNA^Ser^-GCT (**e**) and mito_tRNA^Met^-CAT (**f**) shown as bar plots. Bar colors denote five major cell clusters; expression values represent Z-score normalized abundance.

Beyond nuclear-encoded cytoplasmic tRNAs, we also analyzed mitochondrial tRNAs (mito_tRNAs), which are transcribed independently^65^. Z-score normalized clustering revealed substantial mito_tRNAs expression heterogeneity across hematopoietic cell types (Fig. 4c). Consistent with their metabolic demands, mature lymphoid and myeloid cells exhibited elevated mito_tRNA levels, while quiescent HSCs and early progenitors also displayed high expression of specific mito_tRNAs, in line with their reported high mitochondrial mass despite low respiratory activity (Fig. 4c, d)^65,66^. Moreover, myeloid oligopotent cells exhibited relatively low mito_tRNA isoacceptor expression (Extended Data Fig. 8a), whereas CLPs showed higher levels (Extended Data Fig. 8b). In contrast, lymphoid unipotent cells displayed globally reduced mito_tRNA abundance (Extended Data Fig. 8c). Consistent with this trend, individual mito_tRNA isoacceptors exhibited cell-type-specific enrichment. For example, mito_tRNA^Ser^-GCT was preferentially expressed in CLPs and upstream stem/progenitor populations, whereas mito_tRNA^Met^-CAT was enriched in CLPs and mature lymphoid and myeloid cells (Fig. 4e, f).

Integrating cytoplasmic and mitochondrial tRNA profiles, we identified three distinct isoacceptor subsets with stage-specific expression patterns (Fig. 4c). Subset I, enriched in stem cells and progenitors-may sustain stemness or multipotency. Subset II, predominantly expressed in terminally differentiated cells (e.g., mature lymphoid/myeloid cells), might be associated with mature immune cell state. (Fig. 4c). Subset III showed elevated levels in quiescent or low-proliferation populations, including HSCs, upstream progenitors, and common lymphoid progenitors (Fig. 4c). Previous studies observed the relationship between tRNA subsets and cell proliferation versus differentiation^3^. In our dataset (Fig. 4c), 10 out of 12 tRNAs that were previously classified as “proliferation-associated” cluster within subset II^3^. Conversely, 6 out of 10 tRNAs previously classified as “differentiation-associated” are found in our subset I^3^. These observations suggest that the isoacceptors within each subset may be co-regulated; however, their specific functions in the hematopoietic system still await further investigation.

In summary, our isoacceptor level analysis across 26 continuously differentiating hematopoietic cell types reveals that the cytoplasmic tRNA anticodon pool correlates with cellular differentiation states, potentially reflecting distinct translational programs. Specifically, the cytoplasmic tRNA pool remains largely stable among hematopoietic stem and progenitor cells, while terminally differentiated lineages exhibit compositions distinct from progenitors yet stable within each mature stage. In contrast, mitochondrial tRNAs exhibit greater plasticity; their expression dynamics collectively reflect variations in mitochondrial mass, activity, and functional state.

### tdRs show a preference for highly differentiated cells

While tdRs are increasingly recognized as regulators in RNA silencing and tumorigenesis, their roles in dynamically sculpting cellular identity during differentiation remain enigmatic^2,18,20,21,49,67–70^. To explore the differences in tdRs expression across cell types, we calculated the tdR-to-parental-tRNA ratio by dividing tdR counts by their corresponding parental tRNA isodecoder counts. A clustered heatmap of these ratios revealed that stem cells and multipotent progenitors maintained similarly low tdR abundance, whereas highly differentiated cells displayed markedly higher and more complex patterns of tdR derivation (Fig. 5a and Extended Data Fig. 9a). We observed that 3’tdRs were most abundant in cDCs and pDCs (Fig. 5a). Kernel density estimation (KDE) plots of tdR-to-tRNA ratios highlighted distinct distribution patterns among cell types: upstream progenitors and HSCs showed sharp peaks, indicating in these cells the average tdR ratio is very low, whereas downstream progenitors and mature cells exhibited broader peaks, reflecting both greater variability and a higher overall abundance of tdRs (Fig. 5b and Extended Data Fig. 9b). Notably, downstream cells also presented more right-tailed outliers, demonstrating that specific tRNA isodecoders, such as tRNA^Asp^-GTC-4, tRNA^Cys^-GCA-1, and tRNA^Met^-CAT-1, are more prone to generate tdRs (Extended Data Fig. 10, 11).

**Fig. 5.**
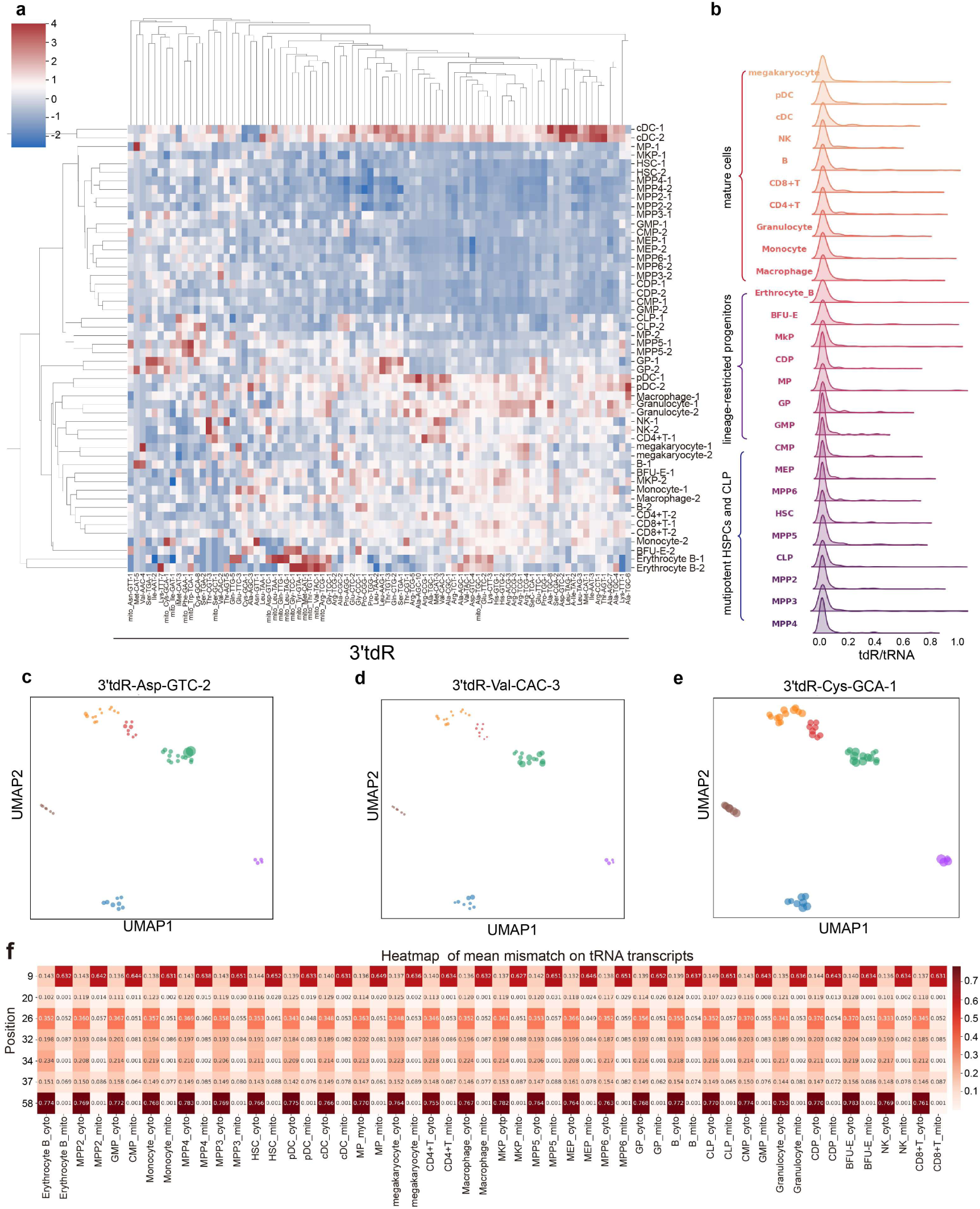
tdR expression and modification patterns of 26 hematopoietic cell types. **a**, Clustered heatmap of 3’tdR expression levels relative to their parental tRNA isodecoders. Data were calculated as the ratio of tdR counts to tRNA isodecoder counts, followed by Z-score transformation for each tRNA isodecoder (clip range -4 and 4). **b,** Kernel density estimation showing the distribution of 3’tdR/tRNA ratios for each unique isodecoder across 26 hematopoietic cell types. **c-e,** UMAP visualization (from Fig. 4d) of 3’tdR expression patterns: 3’tdR-Asp-GTC-2 (**c**), 3’tdR-Val-CAC-3 (**d**), and 3’tdR-Cys-GCA-1 (**e**). Dot size reflects relative expression levels across cell types. **f,** Heatmap of modification abundance at certain sites across 26 hematopoietic cell types (mitochondrial and cytoplasmic tRNAs respectively). Color scale: mean modification levels (two replicates).

We further mapped tdR proportion patterns onto UMAP embeddings generated from tRNA isodecoder expression signatures. The results demonstrated highly diverse tdR patterns among cell types (Fig. 5c**-**e). We found that 3’tdR-Cys-GCA-1 was relatively abundant across all cell types, with only minor variations in its ratio to tRNA (Fig. 5e); however, cDC cells exhibited the lowest expression ratio (Fig. 5a, Extended Data Fig. 10). In contrast, 3’tdR-Asp-GTC-4 (Fig. 5c) and 3’tdR-Val-CAC-3 (Fig. 5d) showed higher proportions in mature lymphoid and myeloid cells (Extended Data Fig. 10). Moreover, 5’tdR-iMet-CAT-3 is abundant in several cell types (Extended Data Fig. 11). These tdRs also displayed greater variability, with the least abundant cell type exhibiting expression levels 100 times lower than the most abundant (Extended Data Fig. 10, 11). Collectively, our findings indicate that tdR expression exhibits cell type-specific variation, with a preferential increase in downstream hematopoietic cells and distinct 3’tdR/5’tdR expression patterns.

### Cell-type– and isodecoder-specific tRNA modification dynamics emerge during differentiation

We also investigated whether tRNA modifications change during mouse hematopoiesis by calculating the mismatch rates for known modification sites across tRNA isodecoders. First, we calculated the mean mismatch rates for each modified position in 26 cell types, separating mitochondrial and cytoplasmic tRNAs. Overall, mismatch rates remained relatively consistent across most potential modification sites. Despite the overall stability in modification rates, we observed variations of over 20% at specific modifications across different cell types, particularly at position 20 (Fig. 5f and Supplementary Table 5).

Next, we focused on the mismatch rates of individual tRNA isodecoders at each position. Modification at position 20, which is reported to be acp^3^U^71^, has recently garnered significant attention due to its connection with glycosylation^50,51^. We identified this modification in mouse morulae, where it was highly enriched in tRNA^Met^-CAT and tRNA^Ile^-AAT (Extended Data Fig. 5a). However, in hematopoietic samples, we observed that in addition to these two tRNA species, tRNA^Ala^-AGC, tRNA^Lys^-TTT, tRNA^Pro^-TGG and tRNA^Cys^-GCA also had this modification, and at a much higher modification rate (Extended Data Fig. 12a, Supplementary Table 5). Interestingly, the modification level at position 20 varied among different isodecoders of the same anticodon, indicating that cells regulate specific isodecoder’s modifications precisely (Extended Data Fig. 12a).

m^1^A58 was consistent across all cell types, with most isodecoders showing over 80% modification rate (Supplementary Table 5). However, we noted clear differences in modification rates between isodecoders of the same isoacceptor. We observed tRNA^Ala^-AGC-2/3 had a very low m^1^A modification rate at position 58, compared to tRNA^Ala^-AGC-4/5 (Extended Data Fig. 12b and Supplementary Table 5). More notably, the dominant isodecoder tRNA^iMet^-CAT-1 exhibited a low abundance of m^1^A58 modification at around 12%, while its minor isodecoder, tRNA^iMet^-CAT-3 (representing 1-2% of total iMet expression), showed a much higher and more divergent m^1^A modification pattern, ranging from 25% to 60% (Supplementary Table 5). These observations reveal isodecoder-specific differences in modification levels, highlighting heterogeneity among tRNA isodecoders of the same isoacceptor.

We also examined modifications at position 34, which is more variable due to the diversity of nucleotides. Categorizing isodecoders by the base at this position revealed distinct trends: A at position 34 was more likely modified to inosine in most isodecoders (Extended Data Fig. 12c), whereas C showed much lower modification rates (Extended Data Fig. 12d). Notably, the modification levels at this position varied between cell types, reflecting cell-type-specific patterns.

To investigate the potential impact of these complex modification patterns on tRNA processing, we analyzed the correlation between the levels of key modifications and the abundance of their corresponding tdRs. First, we examined the relationship between the expression of Dnmt2 (GEO: GSE109125), a known tRNA m^5^C38 modification enzyme that influences tdR generation^22^, and tdR abundance. Our results showed that *Dnmt2* expression was negatively correlated with both 5’tdR and 3’tdR derived from tRNA^Asp^-GTC-4 (Extended Data Fig. 12e). This observation is consistent with previous reports indicating that Dnmt2 regulates tdR formation^22^. Within the normal hematopoietic system, while no clear correlation was observed between m^1^A58 and tdR generation, our analysis of acp^3^U20 revealed specific associations. The levels of 5’tdR-Ala-AGC-1 (Extended Data Fig. 12f), 5’tdR-Lys-TTT-7 (Extended Data Fig. 12g), and 5’tdR-Pro-TGG-1 (Extended Data Fig. 12h) showed significant positive correlations with the abundance of acp^3^U20 on their respective parental tRNAs. In contrast, no statistically significant correlation was observed for tdR-Ile-AAT-2, tdR-Ile-AAT-3, or tdR-Cys-GCA-8. Surprisingly, 3’tdR-Met-CAT-3 (Extended Data Fig. 12i) showed significant negative correlations with the abundance of acp^3^U. Together, these results reveal a set of isodecoder-specific associations between tRNA modification levels and tdR abundance.

In summary, tRNA modifications exhibit dual specificity, spanning both cell types and isodecoders. Notably, a subset of these modifications is associated with isodecoder-specific differences in tdR abundance, suggesting a potential link between certain tRNA modifications and the biogenesis of some specific tdRs in hematopoietic system.

### FLORA-seq mapping of cell-type-specific tRNA isodecoders for guiding sup-tRNA candidate screening

Our discovery of cell-type-specific tRNA isodecoder expression and functional hierarchies inspired precision engineering of sup-tRNAs. The premature termination codon (PTC) leads to a premature termination of protein translation, resulting in loss-of-function disease phenotypes^72–74^. Nonsense mutations account for approximately 11% of all genetic diseases^75^, making them a predominant category of disease-causing mutations in humans (Fig. 6a). A single base mutation can convert 18 codons into stop codons, affecting 10 amino acids in total^16^. sup-tRNAs are a promising therapeutic solution, as they can decode stop codons into sense amino acids, rescuing truncated peptide synthesis. However, selection of optimal endogenous tRNA templates for engineering sup-tRNAs remains a major challenge, owing to the vast number of tRNAs at the isodecoder level. Individual isoacceptors are often composed of dozens of isodecoders, yet how their cell-type-specific expression levels impact suppressor efficiency has remained largely unexplored. For instance, tRNA^Glu^ has two isoacceptors and over 30 isodecoders, complicating the process of efficiently designing tRNAs to read through PTCs for each isoacceptor affected by nonsense mutations (Fig. 6b).

**Fig. 6.**
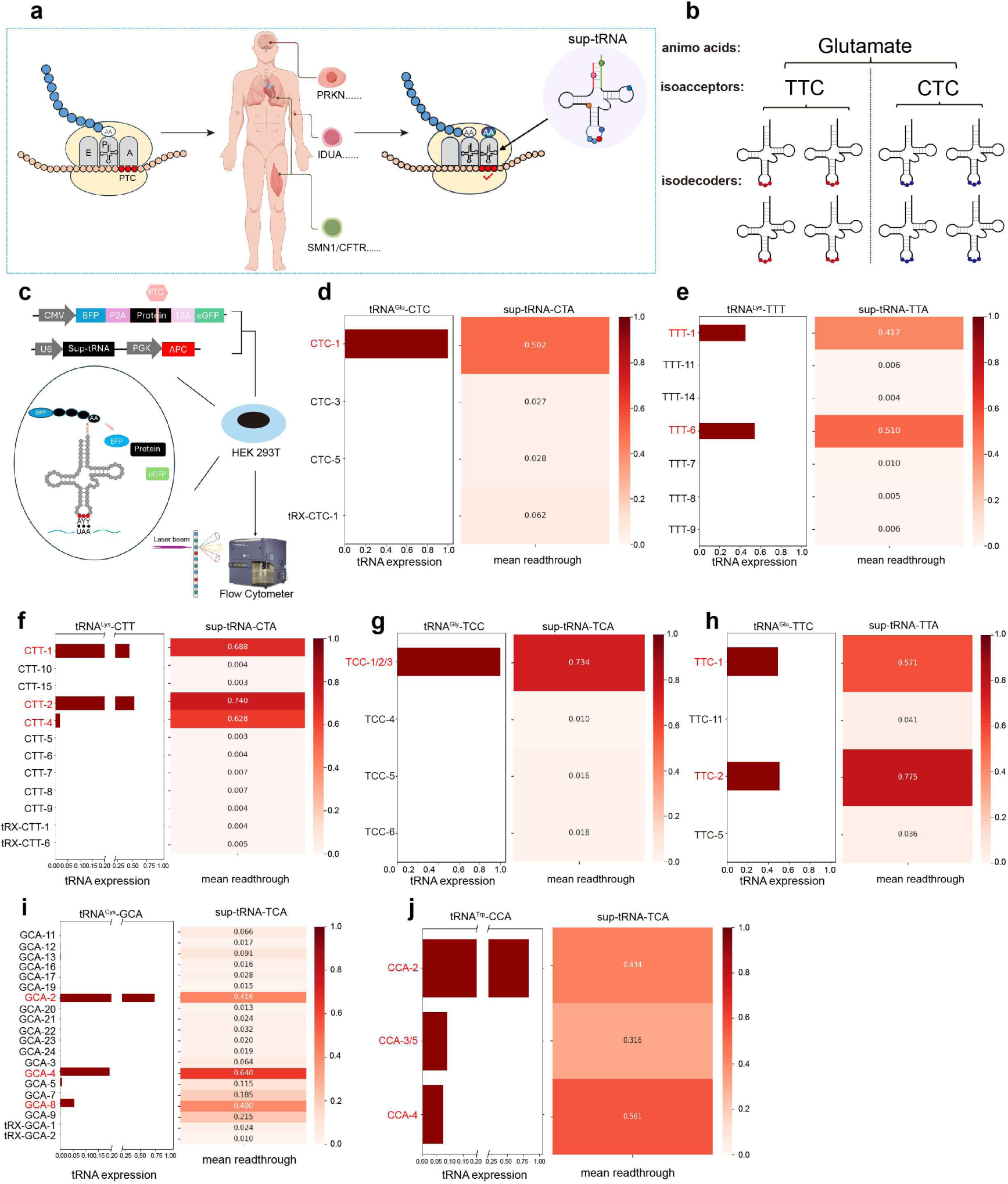
Facilitating suppressor tRNA candidate screening with FLORA-seq. **a**, Schematic of PTC-induced genetic diseases and their rescue by sup-tRNA therapy. Nonsense mutations result in premature termination codons, causing many human diseases. Engineered sup-tRNAs can decode PTCs as sense amino acids. **b,** Schematic illustrating the complexity of tRNA. For example, tRNA^Glu^ has two isoacceptors, tRNA^Glu^-TTC and tRNA^Glu^-CTC, each has many isodecoders. **c,** Reporter system in HEK 293T cells. Its core structure is BFP-P2A– [Test Peptide containing the PTC]-T2A-GFP. GFP expression strictly depends on successful readthrough of the PTC within the test peptide. Two vectors are transfected into cell poll simultaneously. One vector carries CMV promoted florescent protein and PTC containing sequence, the other vector carries U6 promoted sup-tRNA sequence. **d-j,** Expression levels of tRNA isodecoders (left panels) and readthrough efficiencies of their modified sup-tRNA candidates (right panels) for tRNA^Glu^-CTC (**d**), tRNA^Lys^-TTT (**e**), tRNA^Lys^-CTT (**f**), tRNA^Gly^-TCC (**g**), tRNA^Glu^-TTC (**h**), tRNA^Cys^-GCA (**i**), and tRNA^Trp^-CCA (**j**). Expression level: Calculated as the relative percentage of isodecoders within the total tRNA pool. Readthrough rate: Defined as the ratio of GFP-positive to BFP-positive cells, with GFP expression quantified as fluorescence intensity exceeding background threshold. **k,** Schematic diagram of the workflow for screening sup-tRNAs with high readthrough efficiency by FLORA-seq.

To address this challenge, we established a platform in HEK 293T cells to determine whether the expression levels of tRNA isodecoders could provide guidance in designing sup-tRNAs (Fig. 6c). We established a reporter system with two vectors: one carrying a U6 promoted mature sup-tRNA sequence and another carrying two fluorescent proteins flanking a PTC. Successful readthrough of the PTC results in the expression of both blue and green fluorescence, while failure to read through produces only blue fluorescence (Fig. 6c). This system estimates average readthrough rates in cell populations and enables analysis of GFP/BFP ratios at the single-cell level. It also allows high-throughput screening of sup-tRNAs with low background using Flow Cytometry (Fig. 6c).

Using this system, we systematically investigated the relationship between the expression levels of 54 isodecoders and the readthrough efficiencies of their corresponding sup-tRNAs (Supplementary Table 6). For tRNA^Glu^-CTC, the dominant isodecoder tRNA^Glu^-CTC-1 achieved a readthrough rate of 50.2%, while the other engineered tRNAs from rarely expressed isodecoders (less than 1%) can hardly read through PTC, their readthrough rates are below 1% (Fig. 6d). The corresponding sup-tRNAs of major isodecoders tRNA^Lys^-TTT-1 and tRNA^Lys^-TTT-6 exhibited readthrough rates of 41.7% and 51%, respectively (Fig. 6e). The corresponding sup-tRNAs of dominant isodecoders tRNA^Lys^-CTT-1 and tRNA^Lys^-CTT-2 achieved readthrough efficiencies of 68.8% and 74% (Fig. 6f). The tRNA^Gly^-TCC-1/2/3 collectively exhibited an average readthrough rate of 73.4% (Fig. 6g). For tRNA^Glu^-TTC, the major isodecoders demonstrated efficiencies of 57.1% and 77.5% (Fig. 6h). The highly expressed tRNA^Cys^-GCA-2, tRNA^Cys^-GCA-4, and tRNA^Cys^-GCA-8 isodecoders showed readthrough efficiencies of 41.6%, 64%, and 40%, respectively (Fig. 6i). The major isodecoders tRNA^Trp^-CCA-2 and tRNA^Trp^-CCA-4 exhibited readthrough efficiencies of 43.4% and 56.1% (Fig. 6j). Overall, these results indicate that high-efficiency sup-tRNA candidates generally originate from highly expressed cognate isodecoders. Importantly, low-expressed isodecoders (less than 1%) are unlikely to yield high-efficiency sup-tRNAs simply by modifying the anticodon loop.

We next sought to apply this principle in a proof-of-concept setting. Given that our foundational FLORA-seq atlas profiled 26 hematopoietic cell types, we logically focused on validating the platform’s utility in hematologic malignancies driven by nonsense mutations. We selected the MOLT-4 acute lymphoblastic leukemia cell line, which carries a nonsense mutation in the *TP53* tumor suppressor gene. This mutation converts the CGA codon (encoding arginine at position 306) to a TGA stop codon (R306*), thereby introducing the PTC that leads to loss of p53 function and contributes to the cell’s malignant phenotype^76^. Thus, MOLT-4 provides a relevant model for testing a sup-tRNA-based candidate selection strategy. We first performed tRNA sequencing in MOLT-4 cells to identify candidate template isodecoders for engineering a sup-tRNA targeting the TCA (stop) codon. Among the six tRNA^Arg^-TCG isodecoders, only tRNA^Arg^-TCG-1 was highly expressed, with tRNA^Arg^-TCG-2 showing moderate expression (Fig. 7a). We engineered sup-tRNAs from all six isodecoders and transfected them into MOLT-4 cells. Western blot analysis confirmed that sup-tRNA-TCA-1, derived from the dominant isodecoder, restored p53 protein expression most effectively, while sup-tRNA-TCA-2 provided a minor rescue (Fig. 7b). We then assessed the *in vivo* efficacy of sup-tRNA-TCA-1 as a proof of principle. MOLT-4 cells transfected with either this sup-tRNA or a negative control (NC) were intravenously injected into NSG mice to establish a disseminated leukemia model (Fig. 7c). The sup-tRNA pretreatment conferred a significant survival benefit, extending the lifespan of leukemic mice compared with the NC group (Fig. 7d). Body weights remained stable in both groups throughout the study (Fig. 7e). Consistent with improved survival, sup-tRNA pretreatment markedly suppressed leukemia progression. Flow cytometric analysis showed a substantially lower proportion of human CD45 positive MOLT-4 cells in the peripheral blood of the sup-tRNA group at four weeks post-transplantation (Fig. 7f). Longitudinal weekly monitoring further demonstrated that the increase in peripheral leukemia burden over time was significantly attenuated in mice receiving sup-tRNA-pretreated cells (Fig. 7g). Together, these results demonstrate that our expression-level-guided engineering strategy successfully generates a functional sup-tRNA capable of rescuing a disease-relevant nonsense mutation, suppressing tumor progression, and prolonging survival *in vivo*.

**Fig. 7.**
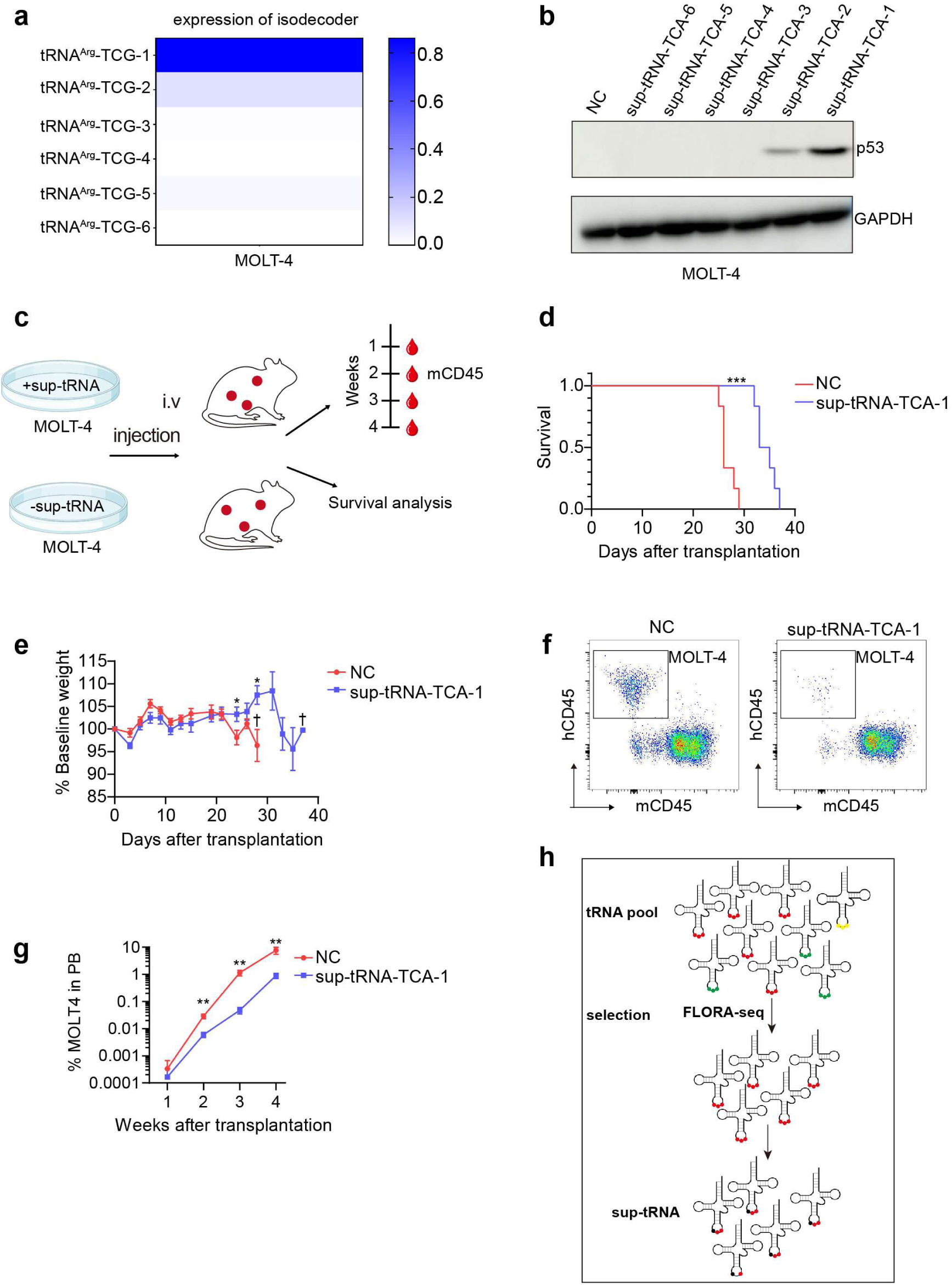
Expression-guided sup-tRNA candidate shows functional rescue in a MOLT4 xenograft model. **a**, Expression levels of the endogenous tRNA^Arg^-TCG isodecoders in MOLT-4 cells by FLORA-seq. **b,** Western blot analysis of p53 protein levels in MOLT-4 cells after transient overexpression of six distinct sequencing-designed suppressor tRNAs (sup-tRNA-TCA-1 to sup-tRNA-TCA-6). GAPDH serves as a loading control. **c,** Schematic of the experimental design: MOLT-4 cells were first transfected *in vitro* with either a negative control (NC) or the candidate sup-tRNA. Subsequently, these pretreated cells were intravenously injected into NSG mice to establish a disseminated leukemia model. **d,** Kaplan-Meier survival curve of NSG recipient mice transplanted with MOLT-4 leukemia cells. **e,** Body weight of mice during the study, expressed as percentage of starting mass. Bars represent mean with SEM. **f,** Representative flow cytometry plots showing identification of MOLT4 (hCD45^+^mCD45^-^) in peripheral blood at 4 weeks post-transplantation. **g,** Percentage of MOLT-4 cells in peripheral blood at the indicated time points. Bars represent mean with SEM. Six mice were included per group. Kaplan-Meier survival analysis (**d**) and two-tailed unpaired Student’s t-tests (**e**, **g**) were used for statistical analysis. \**p* < 0.05, \*\**p* < 0.01, \*\*\**p* < 0.001. **h,** Schematic diagram of the workflow for screening sup-tRNAs with high readthrough efficiency by FLORA-seq.

In summary, our study establishes that endogenous tRNA expression level serves as a key criterion for selecting optimal templates in sup-tRNA engineering (Fig. 7h). Applying this principle, we designed a functional sup-tRNA that rescued p53 expression in a leukemia cell line and significantly delayed disease progression while extending survival in a corresponding mouse model. These findings illustrate how endogenous expression profiles can guide the prioritization of sup-tRNA candidates in a proof-of-principle xenograft model.

## Discussion

tRNA expression and modification landscapes are dynamically regulated under various cellular environments and diseases^1,77^. However, the profiling of tRNAs has been challenging, especially during transitional processes like cell differentiation, due to the large RNA input. Building on the strengths of some pioneering methods^31–44^, we established FLORA-seq to simultaneously detect tRNAs, tdRs, and their modifications from ultra-low RNA inputs. This level of sensitivity approaches that of single-cell analysis, enabling tRNA profiling in critical biological contexts such as embryonic development and differentiation.

Our findings demonstrate that tRNA expression profiles exhibit cell-type specificity and that tRNA isodecoders can be used as markers for cell clustering within a continuous differentiation system. We observed dynamic regulation of tRNA isodecoders expression during hematopoiesis, aligning with the diverse cellular functions at each stage of differentiation. We identify at least 50 high-variance tRNA isodecoders, whose population-associated expression patterns highlight a previously underappreciated layer of tRNA expression heterogeneity. While these patterns suggest potential relevance to differentiation-associated processes, the underlying mechanisms remain to be fully elucidated. This heterogeneity further supports the potential of tRNA-derived features as candidate biomarkers for cell-type classification. Combined with techniques like high-resolution Ribo-seq, single-cell tRNA profiling could provide new perspectives on how codon biased translation regulates lineage commitment and differentiation. Additionally, we observed significant heterogeneity in the expression of tdRs across different cell types, highlighting the critical role of regulating the various aspects of tRNA biology for proper cellular function. Beyond tRNA expression dynamics in hematopoiesis, we observed rearrangement of tRNA modifications across different cell types. Some tRNA modifications are potentially associated with the biogenesis of tdRs derived from a subset of isodecoders, underscoring the distinct roles of individual isodecoders during cell differentiation.

In this study, tRNA profiling across 26 continuously differentiating cell types revealed that changes in anticodon pool composition show a progressive association with distinct cellular differentiation states. The cytoplasmic tRNA anticodon pool tends to remain relatively stable within similar differentiation stages, such as within HSPCs or within terminally differentiated lineages. In contrast, markable differences in anticodon pool composition were observed between these distinct differentiation states. When differentiation state differs significantly (e.g., between stem cells and terminally differentiated cells), the tRNA anticodon pool undergoes more pronounced systematic rearrangement. This aligns with the phenomenon of “tRNA anticodon pool switching between proliferation and differentiation programs” described by Gingold et al.^3^ and may provide a translational adaptive basis for cellular state transitions. Mitochondrial tRNAs displayed greater expression plasticity. Elevated levels of mitochondrial tRNAs were evident not only in terminally differentiated cells, consistent with their increased translational demand, but also surprisingly in HSCs. This observation aligns with previous reports describing a relatively high mitochondrial mass in HSCs despite their quiescent state and low mitochondrial activity^63^. Together, these findings suggest that mitochondrial tRNA abundance in early hematopoietic cells is not strictly coupled to mitochondrial metabolic output, but instead reflects a distinct layer of mitochondrial regulation during early hematopoiesis.

Recent work employing ChIP-seq and tRNA-seq has revealed tRNA expression patterns in distinct settings, including cell-type-specific heterogeneity linked to neuronal vulnerability^11^ and anticodon pool buffering during *in vitro* differentiation^12^. By directly quantifying mature tRNA abundance from ultra-low inputs across a continuous *in vivo* hematopoietic hierarchy, our results demonstrate that isodecoder expression can recapitulate the hematopoietic hierarchy and distinguish cellular identities along differentiation trajectory. Moreover, by simultaneously profiling tdRs and modifications, our approach provides a more comprehensive view of the tRNAome.

The cell-type-specific expression of tRNA isodecoders can further facilitate candidate screening in designing engineered tRNA for PTC diseases. Studies have shown that the efficiency in reading through PTCs of sup-tRNAs derived from different isodecoders can vary substantially^8^. Consistent with a recent study^78^, our analysis suggests that the readthrough efficiency of a sup-tRNA appears to correlate with the endogenous expression level of its parental isodecoder. As recently demonstrated with an AAV-delivered UGA sup-tRNA, tissue-specific differences in sup-tRNA accumulation and aminoacylation correlate with enhanced readthrough in muscle^79^, reinforcing the notion that the native expression pattern of the parental isodecoder can guide sup-tRNA candidate selection. Such differences are likely influenced by isodecoder-specific sequence features that affect functional interactions with the translational machinery, including ribosome binding and competition with release factors, rather than simply differences in transcription efficiency. In current tRNA therapeutic strategies, strong promoters such as U6 are routinely employed to drive the transcription of engineered tRNAs. All sup-tRNAs in our experiments were robustly expressed under the U6 promoter, reaching levels 20–30-fold higher than those of endogenous tRNAs (Supplementary Table 6). However, even under the control of the strong U6 promoter, endogenously silent isodecoders remain challenging to express at high levels in cells. This indicates that elevated transcription alone is insufficient to ensure equivalent functional output across different isodecoders. These observations suggest that the effective cellular concentration of a mature and functional sup-tRNA is shaped not only by promoter-driven transcription, but also by sequence-dependent properties influencing post-transcriptional processing, stability, and aminoacylation. Consistently, we observed that endogenous isodecoders with higher native expression levels tended to exhibit greater PTC readthrough efficiency when engineered as sup-tRNAs. This association suggests that highly expressed endogenous isodecoders may possess intrinsic properties that enhance tRNA functionality and its effectiveness as a sup-tRNA. While the underlying mechanisms remain to be elucidated, our findings provide a practical framework for prioritizing candidate sup-tRNAs based on native isodecoder expression profiles.

Although we can clearly identify a subset of tRNA modifications, as with other sequencing-based approaches, we cannot distinguish between different modifications occurring at the same nucleotide position. In addition, certain modifications do not induce RT stop or misincorporations and therefore remain undetectable by our method. Recent studies have made significant progress toward the simultaneous detection of multiple RNA modifications. For example, Pan-Mod-Seq enables the inference of more than a dozen distinct RNA modifications across many samples in parallel, providing a comprehensive view of ribosomal RNA modification landscapes across species and conditions^80^. Nanopore-based direct RNA sequencing methods have also been developed to simultaneously capture tRNA abundance and modification information from native molecules^44, 81–83^. Beyond fundamental biology, cell-free tRNA modification signatures show promise as diagnostic biomarkers for early-stage colorectal cancer^84^. Future efforts aimed at expanding the repertoire of detectable modifications will further enhance the utility of tRNA sequencing for both fundamental and translational applications.

In conclusion, we established FLORA-seq for ultra-low-input profiling of tRNAs and tdRs, uncover dynamic isodecoder regulation during continuous hematopoietic differentiation, and demonstrate that endogenous isodecoder expression profiles can guide the rational selection of sup-tRNAs.

## Acknowledgments

This work is supported by National Natural Science Foundation of China [82450109, 92581111 to R-J. L; 32501147 to H. L.] and National Key Research and Development Program of China [2021YFA1100800 to R-J. L]; This work is also supported by the Shanghai Frontiers Science Center for Biomacromolecules and Precision Medicine in ShanghaiTech University.

We thank the Molecular and Cell Biology Core Facility (MCBCF), the Multi-Omics Core Facility (MOCF), and the Molecular Imaging Core Facility (MICF) at the School of Life Science and Technology, ShanghaiTech University for providing technical support.

We acknowledge Figdraw (www.figdraw.com) for providing graphical elements used in Fig. 3a and 6a.

## Methods

### Mice

Male C57BL/6J mice (8-10 weeks old) were obtained from Shanghai Jihui Laboratory Animal Care Co., Ltd. Female C57BL/6J mice (8-10 weeks old) were obtained from Shanghai GemPharmatech Co. Ltd. All mouse studies were carried out according to the guidelines of the Institutional Animal Care and Use Committee of ShanghaiTech University.

The *Trmt6*^flox/flox^ mice were kindly provided by Dr. Hua-Bing Li (Shanghai Jiao Tong University School of Medicine).

### Cell culture and transfection

SH-SY5Y, HEK 293T and C2C12 cells were maintained in Dulbecco’s modified Eagle Medium (DMEM) (Corning, 10-013-CV) supplemented with 10% (v/v) fetal bovine serum (Gibco, 10099141C) and antibiotics (penicillin-streptomycin, Meilunbio, MA0110) at 37°C with 5% CO_2_. HEK 293T cells were transfected using EZ trans (Life-ilab, AC04L091).

### Isolation of a given tRNA by biotinylated DNA probes

Total RNA was extracted from HEK 293T cells using TRIzol (Invitrogen). The endogenous tRNAs used in this study were isolated by their own biotinylated DNA probe and were purified by Streptavidin Agarose Resin (Thermo Fisher Scientific). 30 μL of high-capacity streptavidin-conjugated agarose beads were washed with buffer A (10 mM Tris-HCl, pH 7.5) and suspended in buffer B (100 mM Tris-HCl, pH 7.5). Subsequently, 200 μM biotinylated oligonucleotides were mixed and incubated at room temperature for 120 min. After the incubation, the oligonucleotide-coated beads were then washed three times in buffer A and equilibrated in 6 × NTE solution (1× NTE: 200 mM NaCl, 5 mM Tris-HCl, pH 7.5, 2.5 mM EDTA). The oligonucleotide-coated beads and total RNAs in 6 × NTE solutions were heated for 5 min at 70°C and cooled down to 25°C. Then, the beads were washed with 3 × NTE for three times and 1 × NTE for twice. The specific tRNA retained on the beads was eluted with 0.1 × NTE at 70°C and precipitated using 75% ethanol.

Probe of human tRNA^Phe^-GAA: 5’biotin-CTAATGCTCTCCCAGCTGAGCTATTT-3’ Probe of mouse tRNA^Gly^-GCC: 5’biotin-GCATTGGTGGTTCAGTGGTAGAA-3’ Probe of mouse tRNA^Gly^-CCC: 5’biotin-GCATTGGTGGTTCAATGGTAGA-3’

### *In vitro* transcription of tRNA

The tRNA genes were inserted into the pTrc99b vector to construct the pTrc99b-T7-tRNAs plasmid. All tRNAs were synthesized via *in vitro* transcription using T7 RNA polymerase, following previously established protocols^82^. The transcribed tRNAs were then separated using urea-denaturing 12% polyacrylamide gel electrophoresis (PAGE), eluted with 0.5 M sodium acetate (pH 5.2), precipitated with three volumes of ethanol at -20°C, and subsequently dissolved in 5 mM MgCl_2_. To ensure proper folding, the tRNAs were annealed at 85°C for 5 minutes and allowed to cool naturally to room temperature.

Human-tRNA^Phe^-GAA: GCCAAAAUAGCUCAGCUGGGAGAGCAUUAGACUGAAGAUCUAAAGGUCUCUGGUUUGAUCCUGGGUUUCAGAA

### Reverse transcription and RT-qPCR

The purified tRNA and *in vitro* synthetic tRNA were reverse-transcribed using specific primers (Supplementary Table S1) by RT1306, MarathonRT, SuperScript IV (Thermo Fisher) and Maxima H minus (Thermo Fisher) and used for RT-qPCR using a miScript SYBR Green PCR kit (Qiagen). To optimize the temperature and incubation time for reverse RT. Maxima RT at two temperatures (50°C and 60°C) and three RT times (2, 3, and 12 hours). TaqMan Real-Time PCR Assays were performed using reagent kits from Thermo Fisher Scientific, with probes synthesized by Suzhou SYNBIO Co., Ltd.

### RNA mass spectrometry analysis of cellular tRNA modifications

100 ng of specific tRNAs purified by the biotinylated DNA probes was digested with 0.3 μl benzonase, 0.3 μl phosphodiesterase I, and 0.3 μl bacterial alkaline phosphatase in a 20 μl solution including 4 mM NH_4_OAc at 37°C overnight. After complete hydrolysis, 1 μl of the solution was applied to ultra-performance liquid chromatography-mass spectrometry/mass spectrometry (UPLC-MS/MS). The nucleosides were separated on an Atlantis HILIC Silica column (3 μm, 2.1 × 150 mm) and then detected by a mass spectrometer (AB Sciex QTRAP 6500+) in the positive ion multiple reaction monitoring (MRM) mode.

### Ligation of the 3’ OH of RNA to the 5’ pre-adenylated DNA

RNA was heated at 85℃ for 2 minutes and then immediately placed on ice for 2 minutes. The ligation reaction was performed in a 20 µL reaction containing 1 × T4 RNA ligase buffer, 200 ng tRNA, 10 µM pre-adenylated DNA adapter (Synthesized by Novazyme), and 200 U of T4 RNA ligase 2 (NEB) at 25°C for 2 h

### *In vitro* transcription of *Cre* mRNA

*Cre* mRNA was synthesized through *in vitro* transcription from DNA template containing a T7 promoter, utilizing the mMESSAGE mMACHINE T7 Ultra Transcription Kit (Ambion, Thermo Fisher) according to the manufacturer’s protocol. The synthesized *Cre* mRNA was diluted to a final concentration of 100 ng/μl for microinjection.

### Isolation and microinjection of mouse MII Oocytes

To obtain MII oocytes, wild-type (WT) and *Trmt6*^flox/flox^ female mice (6-8 weeks old) were superovulated by injection with 5 IU each of pregnant mare serum gonadotropin (PMSG) (San-Sheng Pharmaceutical), followed by injection with 7 IU of human chorionic gonadotropin (hCG) (San-Sheng Pharmaceutical) 48 h later. MII oocytes were collected 12 h after hCG injection and maintained in HCZB medium for subsequent microinjection. Cytoplasmic microinjection was performed using *Cre* mRNA solution at a concentration of 100 ng/μl.

### *In vitro* fertilization and post-culture sample collection

*In vitro* fertilization (IVF) proceeded immediately after *Cre* mRNA injection and capacitation of sperm from *Trmt6*^flox/flox^ male mice for 20 min in G-IVF PLUS medium (Vitrolife). IVF was conducted in IVF-PLUS medium for 4 h incubated at 37℃ in a humidified atmosphere of 5% CO_2_ and 95% air for 4 hours. After 4 h of IVF, excessive sperm were removed and the fertilized embryos were cultured in G1-PLUS medium (Vitrolife) at 37°C under 5% CO_2_ in air. The embryos at morula stage were lysed in RNAiso Plus reagent (Takara Bio) for RNA extraction.

### Sample preparation and fluorescence-activated cell sorting (FACS)

To ensure sufficient cell numbers, each biological replicate was generated by pooling cells from 2 individual mice. To avoid unrelated variables introduced by batch effect, the collection of all cell populations was finished in a single batch of flow cytometry cell sorting.

Bone marrow (BM) cells were collected by grinding the femurs, tibias, ilia, and spine in FACS buffer (PBS containing 2% fetal bovine serum) in a Petri dish. Prior to the removal of red blood cells by osmotic lysis, 20% of the BM cells were aliquoted for megakaryocyte isolation, and 1% of the BM cells were aliquoted for erythroblast (EryA/EryB) isolation. For megakaryocyte isolation, BM cells were incubated with biotin-conjugated antibodies against Gr1, B220 and CD11b, and megakaryocytes were first enriched by immunomagnetic negative selection using streptavidin magnetic beads (MedChemExpress) before fluorescent staining and sorting. Following the removal of red blood cells, 2.5% of the remaining BM cells were aliquoted for myeloid cell (granulocyte/monocyte/macrophage) isolation and the remainder of the BM cells were used for the enrichment of multipotent HSPCs and committed progenitors. To enrich these lineage^-^ immature cells, the BM cells were incubated with a cocktail of biotin-conjugated lineage antibodies (Gr1, B220, Ter119, CD4, CD8a) and separated by immunomagnetic selection using streptavidin magnetic beads (MedChemExpress). Subsequently, 10% of the lineage-depleted BM cells were aliquoted for erythroid progenitor (BFU-E/CFU-E) isolation, 8% for MkP isolation, 15% for myeloid progenitor (CMP/GMP/MEP/GP/MP/MDP/CDP) isolation, and the remainder for multipotent HSPC (HSC/MPP1-6) and CLP isolation.

Splenocytes were harvested by mashing the spleens through a cell filter (70 µm). After removal of red blood cells by osmotic lysis, 12.5% of the splenocytes were aliquoted for lymphoid cell (B cell/CD4^+^ T cell/ CD8^+^ T cell/NK cell) isolation and the remainder for dendritic cell (pDC/cDC) isolation.

Cells were stained with the optimally titrated fluorochrome-conjugated antibodies for 45 minutes on ice. The combinations of antibodies used for staining are listed in Supplementary Table S2. Dead cells were excluded by DAPI (Beyotime) staining. Cell populations were defined by the combinations of cell surface markers as shown in Supplementary Table S3. All cell populations were sorted on an Aria III with a 70 μm nozzle, except for megakaryocytes, which were sorted on an Aria III (BD Biosciences) with a 100 μm nozzle. The number of cells sorted from the different cell populations is listed in Supplementary Table S3. Flow cytometry data were analyzed using FlowJo software (v10.8.1) (BD Biosciences).

### RNA extraction

Total RNA was extracted from cultured cells and 26 hematopoietic cell types using TRIzol reagent (Invitrogen) according to the manufacturer’s instructions.

For ultra-low-input samples (5 FACS-sorted cells or 5 mouse morulae), total RNA was extracted using the Quick-RNA Microprep Kit (Zymo Research, R1050) following the manufacturer’s protocol.

### Library preparation

#### 1. Deacylation

To remove amino acids from tRNA, deacylation was performed in a 10 µL reaction containing RNA, 150 mM Tris-HCl (pH 9.0) and 10 U of RNase inhibitor (Thermo Fisher) at 37°C for 20 minutes.

#### 2. PNK treatment

To convert the 3’-P or 2’,3’-cP of tRNA into 3’-OH, total RNA was subjected to PNK treatment. The 100 µl reaction comprised 1 × PNK buffer, total RNA, 20 U of T4 PNK (NEB), and 10 U of RNase inhibitor, incubated at 37°C for 30 minutes. RNA purification was performed using the RNA Clean & Concentrator-5 (Zymo Research), and the RNA was eluted with 7 µl of RNase-free water.

#### 3. Poly(A) tailing of RNA using *E. coli* Poly(A) polymerase

RNA was heated at 85℃ for 2 min and then immediately placed on ice for 2 min. The 10 µl reaction comprised denatured RNA, 1 × *E. coli* Poly(A) polymerase reaction buffer (NEB), 1 mM ATP, and 5 U of *E. coli* Poly(A) Polymerase (NEB), then incubated at 37℃ for 5-20 min. The reaction was terminated by heating at 72℃ for 2 min.

#### 4. cDNA synthesis

RNA with poly(A) tails mixed with 2 μL of 5 μM RT primer (Accurate Biology). To anneal RT primer to the template RNA, samples were heated to 72°C for 2 min, then cooled to 25°C. The 25 µL RT reaction comprised 5 μL of RT buffer, 0.5 mM dNTPs, 10 U RNase inhibitor (Thermo Fisher), 10 U/μL Maxima H Minus (Thermo Scientific), 6.25 μL TSO (Accurate Biology) and RNA samples, incubated at 50°C for 2-3 h, then at 85°C for 5 min.

#### 5. Library amplification

To amplify and add adapters to cDNA, 10 μL of 5 × KAPA high GC buffer, 1.5 μL of 10 mM dNTPs, 0.5 μL HiFi (Roche), 2.5 μL i5 index primer (Accurate Biology), 2.5 μL i3 index primer (Accurate Biology) and 8 μL ultrapure water were added 25 μL cDNA and samples were incubated with the following PCR cycling conditions: denaturation (95°C for 3 min), 10-13 × (98°C for 10 s, 70°C for 30 s, 72°C for 30 s), final extension (72°C for 10 min).

#### 6. Purification of library

To remove primer dimers, the cDNA was purified using 1 × DNA Clean Beads (ABclonal). To ensure the proportion of the final effective library, the library was fractionated using a PAGE gel. The bands of 200-500 bp size were excised with a blade and placed into pre-prepared EP tubes. The gel pieces were crushed using a pestle, and the DNA was extracted with low EDTA buffer (10 mM Tris-HCl pH 8.0, 0.1 mM EDTA). After extraction, the DNA was added with 3 volumes of ethanol and 1/10 volume of sodium acetate (Thermo Scientific), along with 1 µL of glycogen, and then precipitated at -80°C.

#### 7. RNA-seq

Each library was sequenced using the TruSeq SBS Kit v4-HS, in paired-end mode with a read length of 2 × 150 bp.

### Plasmid design and construction

The sup-tRNA and reporter constructs were made using standard molecular biology techniques. Details of the constructs are provided as Supplementary Table S1.

### Flow cytometry

Cells were detached by using trypsin (Thermo Scientific, 40124ES60) and collected in 5 mL round bottom polystyrene tube (Falcon, 352054). Cell pellets were washed and resuspended with PBS (Meilunbio, MA0015). Flow cytometry was performed on BD LSRFortessa (BD Biosciences) with channel of BP450/50 (TagBFP), BP515/20 (eGFP), BP780/60 (iRFP670). Data were analyzed with FlowJo (Supplementary Table 6). First, single cells were clustered by FCS and SSC. The cell expressing sup-tRNA were separated by iRFP670. Then readthrough efficiency were calculated by the ratio of cell (BFP^+^GFP^+^) to cell (BFP^+^).

### Preprocessing of sequencing reads

Sequencing reads were preprocessed using Trim_Galore (v0.6.10), with Cutadapt (v4.4) and FastQC (v0.12.1), for adapter removal, quality control, and poly(A) tail removal. Multiple rounds of adapter trimming were conducted to remove Illumina adapters, customized 5’-end adapters, and poly(A) tails. A minimum quality score of 20 was applied to filter low-quality reads. Forward reads with a minimum length of 20 bp were retained for downstream analysis. The command used was:

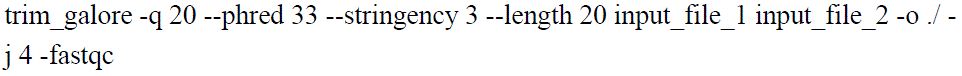

### Alignment and modification identification

tRNA mapping was performed using mim-tRNAseq (v1.3.8). Reads were aligned to human (Hsap) and mouse (Mmus) reference genomes. The initial alignment was conducted with a cluster identity threshold of 0.95 and a maximum mismatch tolerance of 1% of the read length. During realignment, the mismatch tolerance was increased to 5% of the read length. Isodecoders with a coverage threshold below 0.001% of the total reads were filtered out as low coverage. Only uniquely mapped reads were retained for further analysis.

The commands used for each condition are as follows:

mimseq --species Hsap --cluster-id 0.95 --threads 15 --min-cov 0.00001 --max-mismatches 0.075 -n hg38_test --out-dir has_o --max-multi 4 --remap --remap-mismatches 0.05. /tRNA.txt --control-condition HEK 293T

mimseq --species Mmus --cluster-id 0.95 --threads 15 --min-cov 0.00001 --max-mismatches 0.1 -n embryo --out-dir m_emb --max-multi 4 --remap --remap-mismatches 0.05. /tRNA.txt --control-condition wt

mimseq --species Mmus --cluster-id 0.95 --threads 15 --min-cov 0.00001 --max-mismatches 0.1 -n blood --out-dir m_blood --max-multi 4 --remap --remap-mismatches 0.05. /tRNA.txt --control-condition HSC

### Modification detection, read stops, and readthrough rate

Modification signatures and readthrough data were extracted from the mim-tRNAseq results. The proportion of full-length transcripts was defined as the fraction of reads where the mapped length exceeded 95% of the corresponding tRNA transcript length. For each tRNA isodecoder, the total number of full-length and non-full-length reads was counted.

The readthrough rate was calculated as the ratio of read counts at the -1 and +1 positions adjacent to known or potential modification sites. Isodecoder coverage was determined using the sum of read counts for each base of the tRNA isodecoders, calculated with the pysam pileup function. Data were visualized by grouping tRNA transcripts with the same isodecoder or by computing the median and upper/lower quantiles of coverage information for each base position.

### Identification and quantification of tdRs

tdRs were defined as reads mapped to tRNA isodecoders with a mapped length of less than 40 nucleotides. tdRs were separated from uniquely mapped BAM files using pysam (v0.21.0). These reads were further classified into 5’tdRs and 3’tdRs based on their mapped locations: Reads starting and ending at positions < 30 were categorized as 5’tdRs. Reads starting at positions > 30 were categorized as 3’tdRs. Reads starting at positions < 30 and ending at positions> 30 were catagoried as i-tdRs.

Read counts for tdRs were obtained using custom scripts. Differential expression of tdRs between WT and *Trmt6* knockout (KO) mouse embryos was analyzed with pyDESeq2 (v0.4.9) using two replicates for each condition. Differential analysis was conducted at both the isodecoder level and the isoacceptor level separately.

### Blood system tRNA distribution analysis

Data cleaning and mapping were performed as described for HEK 293T and embryo samples. An AnnData object was created using scanpy (v1.9.8) for further analysis. tRNA isodecoders were filtered based on their presence in at least two samples. Sample sizes were normalized to 10,000 reads. Relative expression dot plots were generated with scanpy. For clustered heatmaps, expression data were further normalized within the same tRNA isodecoder across all cell types using scipy (v1.7.3) Z-scores. Principal component analysis (PCA) and k-nearest neighbors (KNN; n_neighbors = 4, n_pcs = 20) were applied to obtain clustering information, which was then visualized using the UMAP function. Clusters were grouped with the Leiden algorithm. Trajectory inference was performed using PAGA (n_neighbors = 5), and pseudotime was calculated based on the trajectory results, with the HSC group specified as the root point. Relative tRNA isodecoder expression levels along the differentiation path were plotted using normalized data.

For tdRs in the blood system, tdR reads were extracted as previously described. tdR read counts were normalized between samples using the same matrix as for tRNA summed counts sample size normalization. Separate clustered heatmaps were created for 5’tdRs and 3’tdRs.

The tdR/tRNA ratio was calculated within each sample by dividing the total tdR read counts by the corresponding tRNA read counts derived from the same sample.

Clustered heatmaps and kernel density estimation (KDE) plots were generated to illustrate the distribution of levels of tdR generation for different tRNA isodecoders and cell types.

### Plotting

All heatmaps, clustered heatmaps, violin plots, volcano plots, radar plots, bar plots, scatter plots, kernel density estimation (KDE) plots, and isodecoder coverage plots were generated using matplotlib (v3.5.3) and seaborn (v0.12.2). Dot plots, UMAP visualizations, and pseudotime trajectories were created using scanpy (v1.9.8).

**Mean expression:** counts were first normalized between group to a sum of 1e4, then calculated mean of 2 replicates for the same cell type.

**High-variance isodecoder:** High-variable isodecoders were defined based on expression variability using the Scanpy implementation of highly variable gene (HVG) selection (sc.pp.highly_variable_genes). Specifically, isodecoders were first binned according to their mean expression levels. Within each bin, the dispersion (variance-to-mean ratio) of each isodecoder was normalized by subtracting the mean and dividing by the standard deviation of dispersions computed for all isodecoders in that bin. Isodecoders with a normalized dispersion above the threshold (min_disp = 0.3) were classified as high-variable isodecoders.

